# Loss of JAK1 Function Causes G2/M Cell Cycle Defects Vulnerable to Kif18a Inhibition

**DOI:** 10.1101/2025.02.19.638911

**Authors:** Vanessa Kelley, Marta Baro, William Gasperi, Nicholas Ader, Hannah Lea, Hojin Lee, Chatchai Phoomak, Lilian Kabeche, Megan King, Joseph Contessa

## Abstract

To gain insight into biological mechanisms that cause resistance to DNA damage, we performed parallel pooled genetic CRISPR-Cas9 screening for survival in high risk HNSCC subtypes. Surprisingly, and in addition to ATM, DNAPK, and NFKB signaling, JAK1 was identified as a driver of tumor cell radiosensitivity. Knockout of JAK1 in HNSCC increases cell survival by enhancing the DNA damage-induced G2 arrest, and both knockout and JAK1 inhibition with abrocitinib prevent subsequent formation of radiation-induced micronuclei. Loss of JAK1 function does not affect canonical CDK1 signaling but does reduce activation of PLK1 and AURKA, kinases that regulate both G2 and M phase progression. Correspondingly, JAK1 KO was found to cause mitotic defects using both EdU labeling and live cell imaging techniques. Given this insight, we evaluated Kif18a inhibition as an approach to exacerbate mitotic stress and enhance the efficacy of radiation. These studies establish Kif18a inhibition as a novel strategy to counteract therapeutic resistance to DNA damage mediated by G2 cell cycle arrest.

## INTRODUCTION

Radiation is a therapeutic modality utilized for treatment of ∼60% of all cancer patients^1^. For head and neck squamous cell carcinoma (HNSCC), radiation in combination with systemic therapy (cisplatin or cetuximab) is the standard of care for locally advanced disease. While well selected patient populations, such as non-smokers with early-stage HPV+ and p53 intact tumors, can achieve local control rates with radiation that approach 100%, patients with unfavorable risk factors can fail locally or regionally at rates of 50% or more^2^. Identifying mechanisms of therapeutic resistance to radiation and other DNA damaging agents is therefore a critical step for the design and deployment of new and more effective interventions.

Tumor cells evade radiation-induced cell death by reducing formation of reactive oxygen species (ROS) and subsequent DNA damage^3,4^, enhancing DNA repair *per se*^5,6^, up-regulating pro-survival signaling programs (e.g. EGFR), or adapting to cellular or physiologic states that enable these processes to occur (reviewed in Huang, *et al.*^7^). Among these cellular states, regulation of the cell cycle has been recognized as a principal mechanism for enabling DNA repair and enhancing tumor cell survival^8^. DNA damage induces arrests in either G1 or G2 phases. However, in tumor cells with p53 mutation, G2 arrest is the major safeguard for preventing the mitotic failure and cell death induced by unrepaired DNA damage. The most widely studied activation switch for G2 arrest is an increase in CDC25C phosphatase mediated dephosphorylation, relative to kinase dependent inhibitory phosphorylation, of cyclin dependent kinase-1 (CDK1) (reviewed in Hochegger, *et al.*^9^). CDK1 in complex with cyclin B1 phosphorylates numerous substrates to regulate chromatin condensation and progression to mitosis. Because CDK1 inhibitory phosphorylation is primarily regulated by Wee1, this kinase has become an attractive target for mitigating the protective G2 arrest and driving DNA damage-induce mitotic cell death.

In this work, we leverage pooled genome screening using fractionated delivery of ionizing radiation to identify mechanisms of tumor cell resistance in high risk (HPV-, p53 mutant) HNSCC models. This design, with delivery of radiation over several days, allows for the experimental results to be driven by and to unfold in the context of recurring DNA damaging and the cellular responses that resolve damage to ensure survival. The results of these studies demonstrate pathways known to regulate survival after DNA damage, such as DNA damage sensing and repair kinases and NFKB signaling proteins, as well as unanticipated gene targets with heretofore uninvestigated functions in the regulation of cell survival after DNA damage. We discover and demonstrate that JAK1 not only regulates G2 progression, but also M phase progression, revealing a dual role that both explains therapeutic resistance and identifies Kif18a inhibition as a tractable intervention for enhancing mitotic cell death in DNA damage resistant tumors.

## RESULTS

### CRISPR-Cas9 screening identifies mechanisms of radiation sensitivity or resistance

To identify pathways that regulate cell survival after exposure to ionizing radiation (IR), we performed parallel pooled genome CRISPR-Cas9 genetic screens in HPV negative Cal27 and Detroit562 HNSCC cell lines. Because kinases are major drivers of adaptive cell signaling pathways, the Brunello kinome-wide gRNA library (which targets 763 signal transduction related genes^10^) was reconstituted and sequenced to ensure gRNA representation (Supplemental Figure 1a, b). Controls were compared to experimental groups treated with 2Gy/day for a total of four doses to simulate clinical radiation therapy regimens. *A priori* we anticipated enrichment or depletion of gRNAs for kinases that regulate survival through reactive oxygen species formation, DNA damage and repair, genome stability, mitosis, and cell death (Figure 1a). Screening for each cell line was performed with three biologically independent replicates, and gRNA prevalence between irradiated and non-irradiated samples was determined by deep sequencing as previously described^3^. Significant differences for gRNA abundance were determined by MAGeCK analysis^11^, and q values (false discovery rate corrected p values) for the corresponding genes are plotted for both Cal27 (Figure 1b) and Detroit562 (Figure 1c). Significantly depleted gRNAs (shown in red) suggest that loss of function confers radiation sensitivity, and significantly enriched gRNAs (shown in green) suggest that knockout (KO) causes radiation resistance.

**Figure 1:**
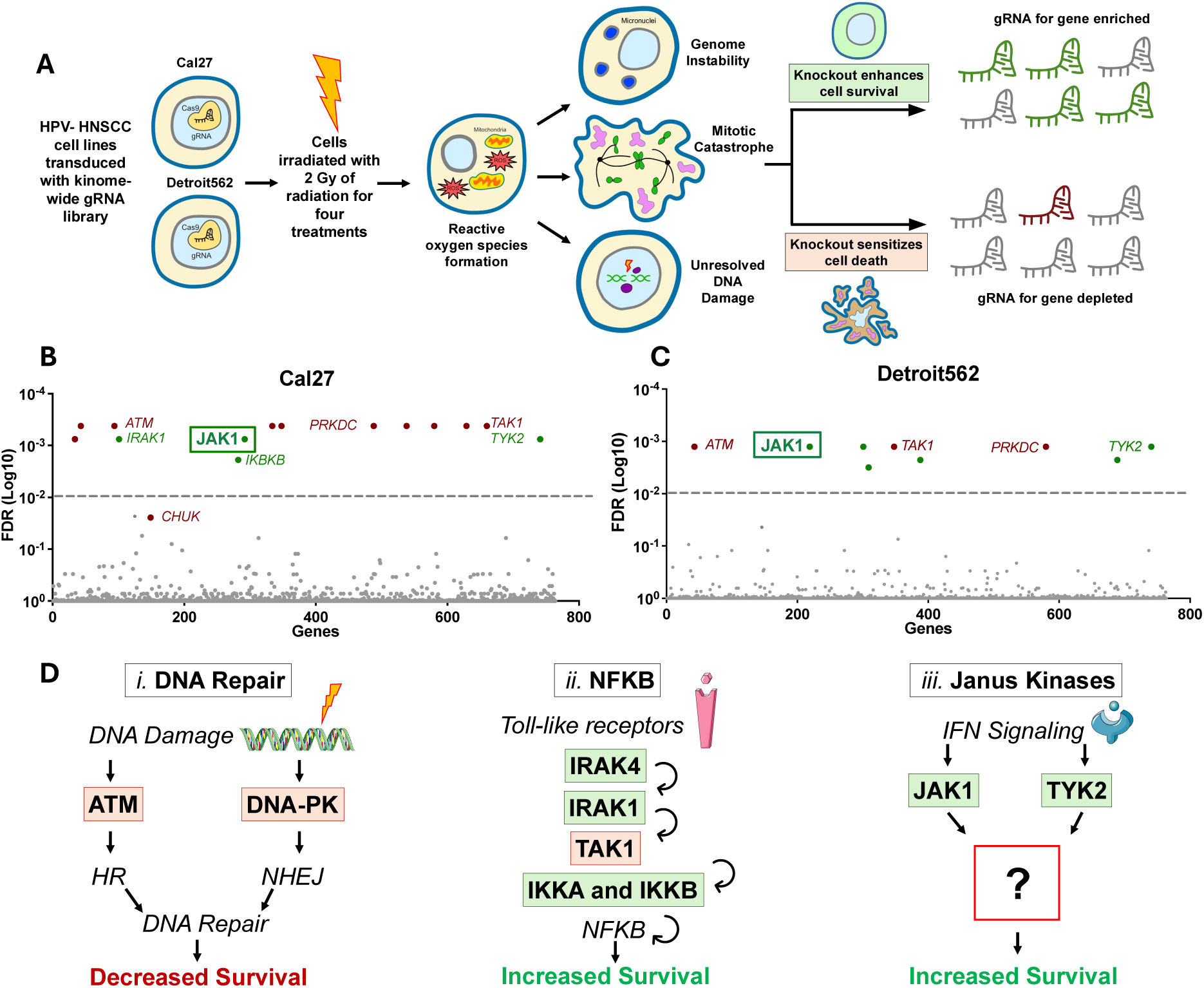
Radiation CRISPR screen in HNSCC cell lines reveals that JAK1 regulates survival after radiation exposure. (**A**) Schematic of radiation CRISPR-Cas9 screening methodology. Cal27 and Detroit562 HSNCC cell lines were engineered to stably express Cas9, transduced with a kinome-wide gRNA library, and irradiated with 2 Gy for four consecutive treatments. Two weeks after treatment, gRNAs were isolated, PCR amplified, and sent for deep sequencing in parallel with unirradiated controls. Simplified screening results are shown for (**B**) Cal27 and (**C**) Detroit562 cell lines. Hits with an FDR ≤ 0.01 were considered significant. (**D**) Summary of pathways identified by the CRISPR screens including (*i)* DNA repair, (*ii*) NFKB signaling, and (*iii*) JAK signaling.

The screening results identified three pathways that alter the cellular radiation response (Figure 1d). Both cell lines demonstrated significant depletion of gRNAs targeting the *ATM* and *PRKDC* genes (Supplemental Table 1a, b; q<0.01) consistent with their known function in DNA damage sensing and repair^12^. Multiple gRNAs for genes in the NFKB signaling pathway were also identified: *IRAK1, IKBKB* (q<0.01), *CHUK* (IKKA, q<0.05), and IRAK4 (which was of borderline significance p=0.0048, q=0.23) (Supplemental Table 1a, b). These enrichments suggest that loss of NFKB signaling enhances radiation resistance^13,14^ (Figure 1d). Guide RNAs targeting *MAP3K7* (also known as TAK1, a kinase that is downstream of IRAK1/4) were significantly depleted in the irradiated group for both the Cal27 and Detroit562 cell lines (q<0.01). We interpret these results to reflect the complex role of TAK1 as an integration point for signaling from NFKB, p38 MAPK, and ATM and reinforce its known role in protection from genotoxic stress^15^. Together, the enrichment of gRNAs for the known DNA repair and NFKB pathway proteins validate the methodology and demonstrate the overall success of this pooled genetic CRISPR-Cas9 screening strategy.

Beyond these known pathways, gRNAs targeting JAK1 and TYK2 were also found to be significantly enriched in both Cal27 and Detroit562 cells following radiation treatment (Figure 1b-c). These results suggest that loss of JAK1 or TYK2 promotes resistance to radiation. JAK1 and TYK2 are both Janus kinase family members that are downstream effectors of Interferon (IFN) and Interleukin (IL) receptor mediated signaling. Theses kinases stimulate inflammatory signaling that can cause either cell growth or differentiation, but have no defined role in directly regulating tumor cell intrinsic responses to radiation. We therefore sought to understand how loss of JAK1 kinase function could enable tumor cell survival after radiation therapy.

### Loss of JAK1 function causes resistance to ionizing radiation

To validate the CRISPR-Cas9 genetic screening results, we generated isogenic JAK1 knockout (KO) cell lines with two different JAK1-targeting gRNAs or a non-targeting gRNA control using CRISPR-Cas9 in each cell type (Supplemental Table 2). After clonal isolation, loss of JAK1 protein expression was confirmed by Western blot (Figure 2a) and genomic sequencing (Supplemental Figure 2a-b). In addition, JAK1 KO blocked or significantly reduced IL-6, IFN-⍺, and IFN-ɣ signaling to STAT1 and STAT3, and suppressed protein levels of the downstream transcriptional target IRF9 in both cell lines (Figure 2d-e).

**Figure 2:**
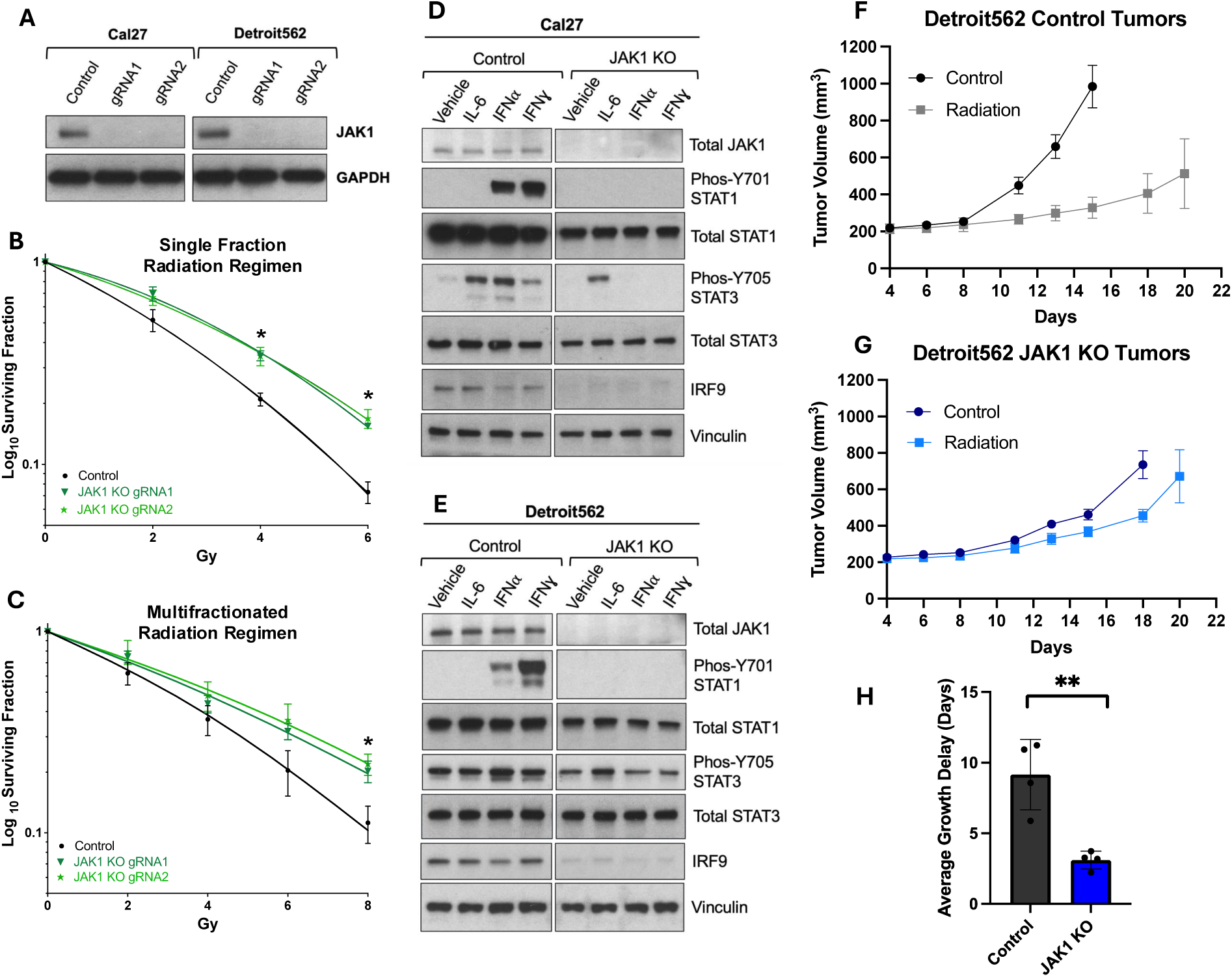
JAK1 knockout causes radioresistance. (**A**) Cal27 and Detroit562 JAK1 KOs were generated with two individual sgRNAs and loss of JAK1 protein was confirmed by western blot. (**B**) Single fraction or (**C**) multiple fraction clonogenic survival results comparing Cal27 control (transduced with a NT gRNA) and JAK1 KO. IFN and IL signaling in (**D**) Cal27 and (**E**) Detroit562 control or JAK1 KO cells stimulated with 50ng/ml of IL-6, 100ng/ml of IFNA, and 4ng/ml of IFNg for one hour. (**F**,**G**) Detroit562 control or JAK1 KO tumor growth kinetics treated with or without fractionated radiation (2Gy/day for 10 days). (**H**) Average tumor growth delay (time to reach 500mm^3^) of control or JAK1 KO tumors receiving radiation treatment. * indicates p< 0.05. ** indicates p<0.01

Cell survival after exposure to radiation was then determined for each isogenic HNSCC pair. In Cal27, *in vitro* clonogenic survival analysis demonstrated enhanced survival for JAK1 KO with statistically significant differences in the range of 4-8Gy for both single and multiple fraction regimens (p<0.05, Figure 2b-c). For Detroit562, the effect of JAK1 KO on radiosensitivity was examined using xenografts because this cell line less readily forms colonies *in vitro* (Figure 2f-h). Following inoculation, both control and JAK1 KOs cells were 100% effective at establishing tumors. However, JAK1 KO tumors progressed at a slower rate. This finding is consistent with results from human melanoma xenografts with JAK1 KO, which also display reduced growth rates in athymic mice^16^. As anticipated, fractionated radiation treatment (2Gy/day) significantly reduced tumor growth in controls by an average of 9 days. However, radiation had only a minor effect on Detroit562 JAK1 KO tumors (Figure 2f) with a non-significant average delay of 3 days. The loss of tumor response to radiation treatment both *in vitro* and *in vivo* unequivocally demonstrates that JAK1 KO promotes radioresistance.

To assess whether transcriptional programs related to DNA damage and repair were altered by JAK1 KO, RNA sequencing of three independent replicates for each isogenic cell line pair was performed (Supplemental Data Files 1 and 2). The results show global downregulation of Janus Kinase-associated genes (Supplemental Figure 2c, f). However, gene set enrichment analysis did not identify a statistical difference in the gene signature associated with DNA repair for either Cal27 or Detroit562 cells (Supplemental Figure 2e, h). This data indicated that modification of transcriptional programs to enhance DNA damage sensing and repair was unlikely to be the mechanism that promotes radiation resistance.

### JAK1 regulates DNA damage-induced micronuclei formation

Interactions of DNA damage and repair pathways with proteins traditionally linked to immune signaling with have garnered significant interest^3,5,17^, and we have recently observed a role for cGAS-STING signaling in promoting radiation-induced DNA damage. We therefore quantified cGAS localization in irradiated control or JAK1 KO cells (Figure 3). In both Cal27 and Detroit562 controls, a robust induction of DAPI-positive micronuclei was observed 24h after treatment with 4Gy (Figure 3b, h), with approximately 80% of these micronuclei staining positive for cGAS, consistent with prior reports^17^. Strikingly, both Cal27 and Detroit562 JAK1 KO cells exhibited a ∼50% decrease in radiation-induced DAPI-positive and cGAS-positive micronuclei (Figure 3e, k). In addition, treatment with abrocitinib, a selective JAK1 inhibitor^18^, showed a dose dependent decrease in micronuclei formation in control cells (Figure 3c, i). At 1uM, abrocitinib reduced radiation-induced micronuclei formation by an average of 33% and 38% in Cal27 and Detroit562 cells, respectively, with similar results observed for cGAS-positive micronuclei. Importantly, and consistent with on-target activity of the inhibitor, there was no effect of abrocitinib on radiation-induced micronuclei formation in JAK1 KO (Figure 3f, l). Together, the consistent reduction of micronuclei formation in each cell type suggests that JAK1 KO or inhibition is associated with an overall enhancement of chromosome stability.

**Figure 3:**
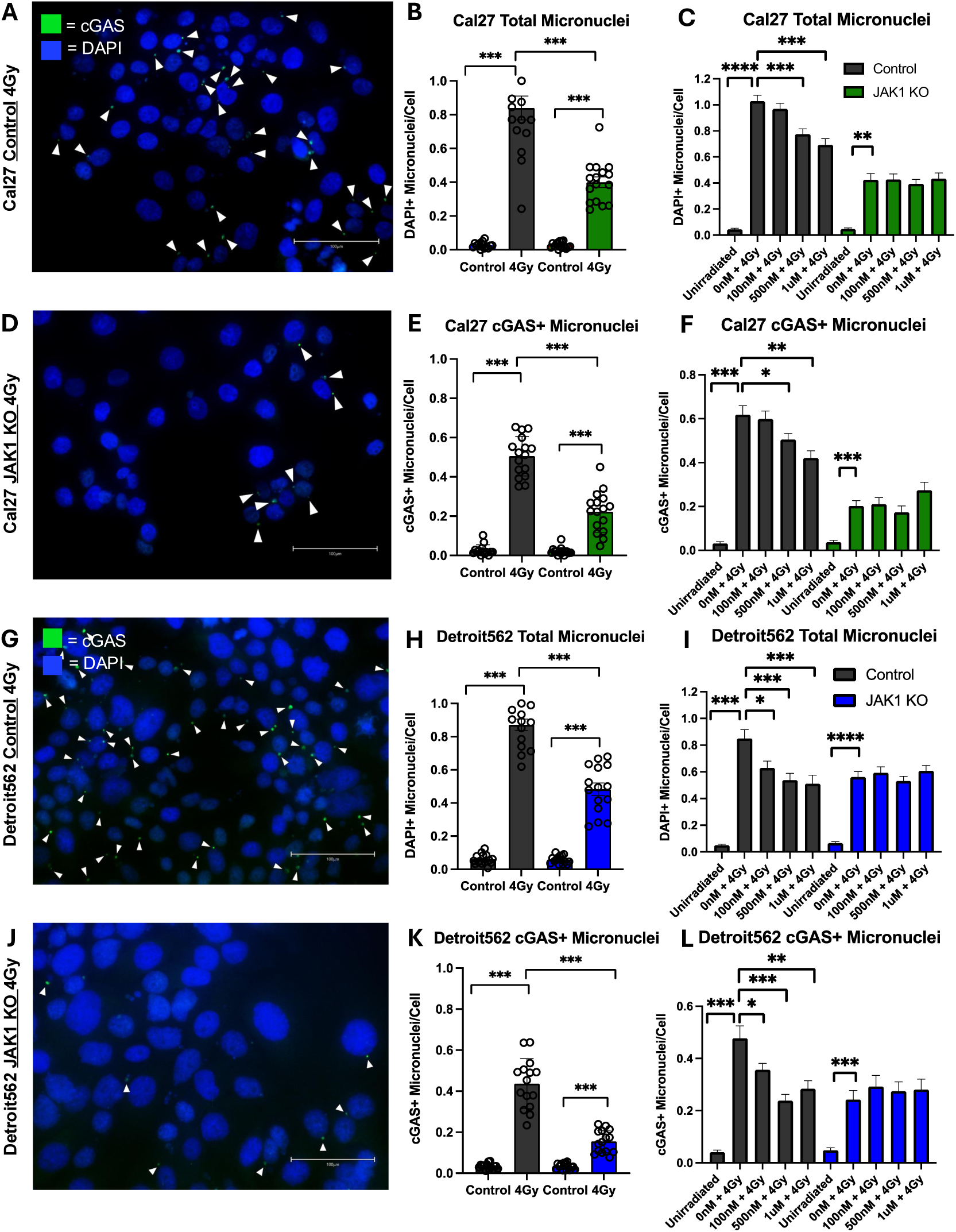
Loss of JAK1 function reduces radiation-induced micronuclei formation. Fluorescence microscopy of (**A**, **D**) Cal27 or (**G**, **J**) Detroit562 control or JAK1 KOs stained for DAPI-positive DNA content and cGAS 24h after treatment with 4Gy. The number of (**B**, **H**) DAPI-positive and (**E**, **K**) cGAS-positive micronuclei were quantified for each field. (**C**, **F**) Cal27 or (**I**, **L**) Detroit562 control and JAK1 KO cells were pre-treated with 0-1uM of abrocitinib for 1 hour and irradiated with 4Gy. DAPI-positive and cGAS-positive micronuclei were then quantified for each field. * indicates p< 0.05. ** indicates p<0.01. *** indicates p<0.005. **** indicates p<0.001.

### JAK1 regulates cell cycle transitions after DNA damage

DNA damage activates a G2 cell cycle arrest, allowing time for DNA repair, but unrepaired DNA damage leads to chromosome mis-segregation events during mitosis with subsequent formation of micronuclei. We therefore investigated the effects of JAK1 KO on cell cycle progression with propidium iodide (PI) labeling 24 hours following exposure to radiation. In Cal27, 8Gy caused an approximate doubling of the G2/M population in JAK1 KO cells compared to controls 24 hours after IR (27% vs 52%; p<.01; Figure 4a-b) with corresponding reductions of both G1 and S populations (Figure 4c-d). In addition, the appearance of an 8N tetraploid population was observed following radiation treatment in JAK1 KOs (Figure 4b, Supplemental Figure 3a). Surprisingly, Detroit562 JAK1 KOs had a significantly increased proportion of cells in the G2/M phase of the cell cycle observed at baseline (20% vs 43%; p<.01; Figure 4e). Although there was no further enhancement of this population after radiation (Figure 4h), a significant increase in tetraploid cells was observed after 8Gy in JAK1 KOs compared to controls (14% vs. 32%; p<.01: Figure 4f, Supplemental Figure 3d). Together, this data indicates that JAK1 KO leads to DNA damage-induced G2/M arrest.

**Figure 4:**
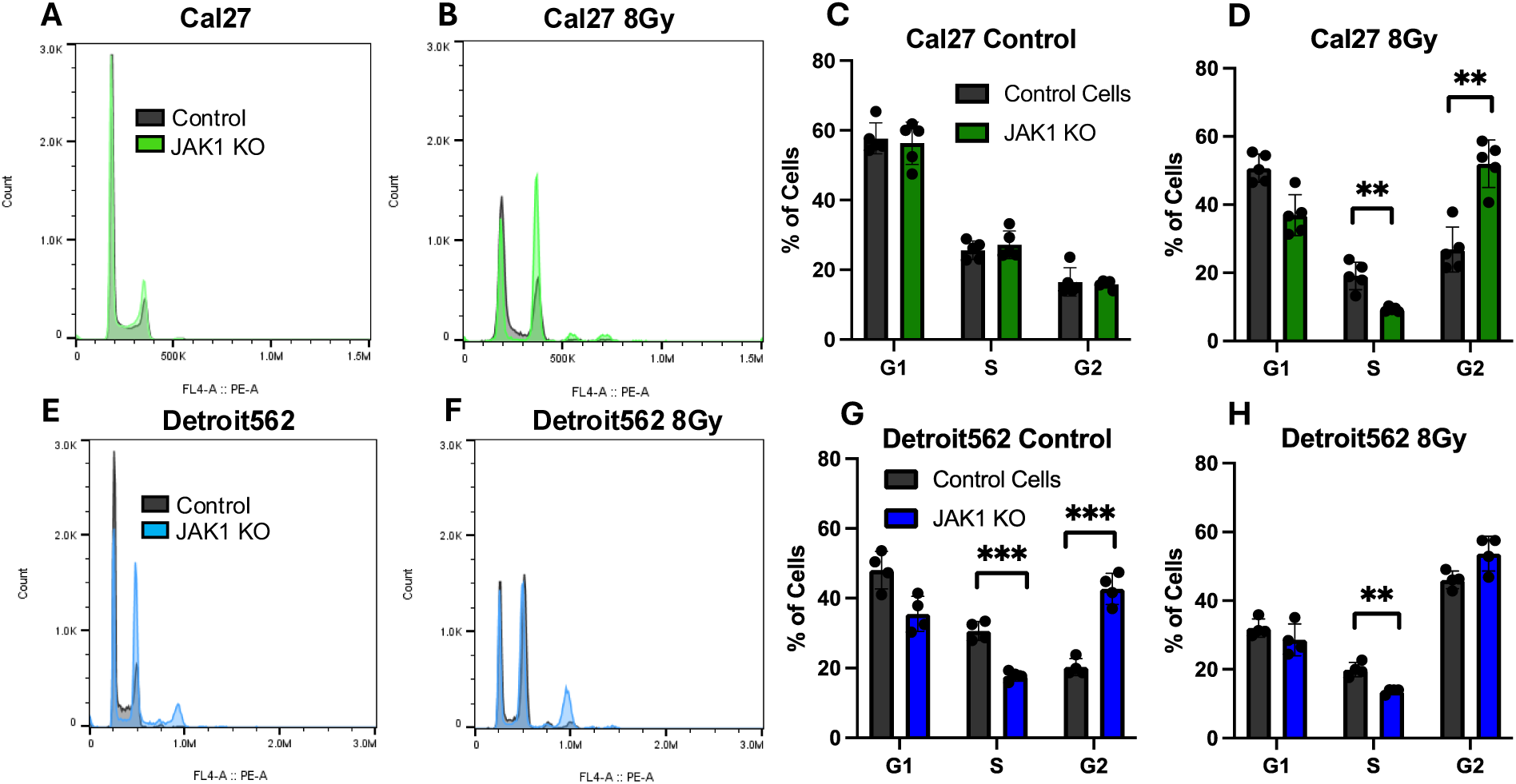
JAK1 KO enhances G2/M cell cycle arrest. Cell cycle profiles of (**A**,**B**) Cal27 or (**C**,**D**) Detroit562 with or without JAK1 KO 24 hours following control or radiation (8 Gy) conditions. Cell cycle phase for control and irradiated cultures was quantified for four independent experiments for either (**C**,**D**) Cal27 or (**G**,**H**) Detroit562. ** indicates p< 0.01. *** indicates p<0.001

Increases in G2/M populations arise from either expedited progression through S phase or from decreased progression through G2 and M phases. To distinguish between these two possibilities, EdU experiments were performed to track an S-phase labeled population. Cal27 and Detroit562 control and JAK1 KO cells were pulsed with EdU, irradiated with 8Gy, and distributions for the EdU-labeled population at 8 and 24 hours after treatment were examined. In Cal27 cells, EdU staining showed initiation of a G2/M arrest at 8 hours following radiation in both control and JAK1 KOs (Supplemental Figure 3b-c). However, in agreement with PI labeling experiments, the G2/M arrest was sustained in cells with JAK1 KO at 24 hours (Figure 5a-b). JAK1 KOs also showed a significantly reduced population of labeled G1 phase cells compared to controls (53% vs 20%, p<.01) indicating a deficiency in G2/M progression. A similar decrease in the EdU-labeled G1 population was found in Detroit562 JAK1 KOs compared to controls, confirming a G2/M progression defect in this cell line (42% vs. 18%, p<0.1; Figure 5e). EdU labeling also revealed a second round of DNA synthesis between the G2 and tetraploid populations detected in Cal27 and Detroit562 cells (Figure 5b, Supplemental Figure 3f). This 8N population was particularly enriched in irradiated Detroit562 JAK1 KOs (8N population 11% vs 44%, p<.01; Figure 5f). Together, these results exclude the possibility that JAK1 KO enables early S phase progression and instead indicate deficiency in G2 as well as M phase abnormalities that lead to 8N tetraploid cells.

**Figure 5:**
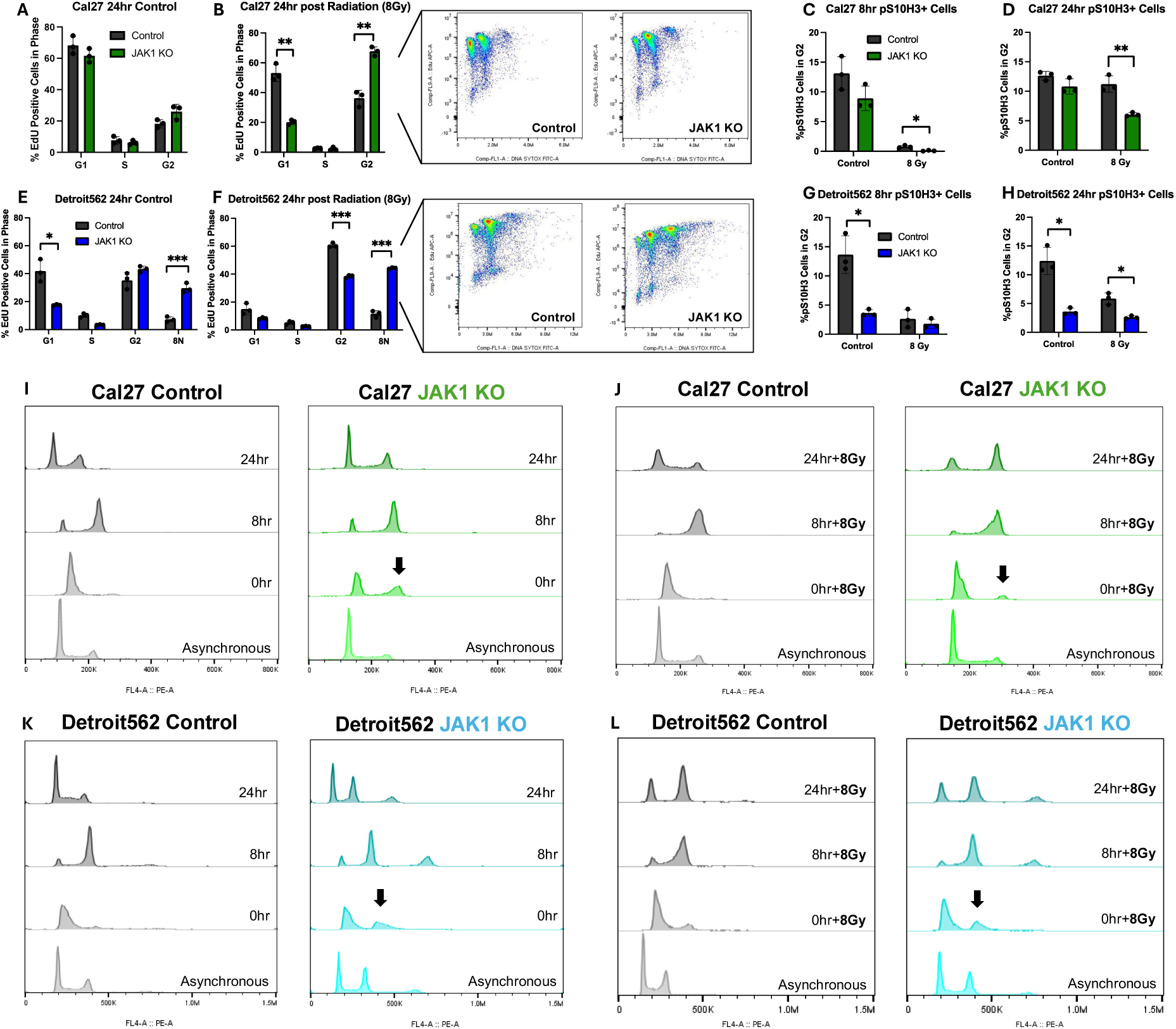
Loss of JAK1 disrupts cell cycle progression. (**A**,**B**) Cal27 control and JAK1 KOs were EdU labeled and irradiated (8Gy) followed by treatment with SYTOX green 24hrs later to determine DNA content distribution. The inset shows distributions for the EdU-labeled population. (**C**,**D**) Phospho-S10 H3 signal was quantified for control or irradiated (8Gy) Cal27 and JAK1 KOs 8- or 24-hours following treatment. The percentage of pS10 H3-positive events in the G2 population is shown for each timepoint. (**E**-**H**) Identical experiments for isogenic control and JAK1 Detroit562 with inclusion of the 8N population. (**I**,**J**) Cell cycle distribution time courses for Cal27 or (**K**,**L**) Detroit562 control or JAK1 KO cells synchronized using a double thymidine block. Cells were harvested at indicated timepoints following release and control or radiation (8Gy) treatment. * indicates p< 0.05. ** indicates p<0.01. *** indicates p<0.005.

The effect of JAK1 KO on phosphorylation of Histone H3 at serine S10 (pS10-H3), a marker for condensed mitotic chromatin^19^, was also examined after radiation treatment. At 8 hrs Cal27 JAK1 KOs displayed significantly less pS10-H3 signal in the G2/M population with ∼10-fold fewer positive cells (Figure 5c). This reduction in chromatin condensation was also sustained, with significantly fewer pS10H3 positive cells at 24hrs (11% vs 6%, p<.01; Figure 5d). Detroit562 JAK1 KO cells also showed differences at baseline with 4% pS10 H3 positivity in G2/M vs. 14% in controls (p<.05; Figure 5g), as well as significantly fewer pS10-H3 positive cells 24hrs after radiation (3% vs 6%; p<.05: Figure 5h). Interestingly, pS10-H3 signal was increased in tetraploid Detroit562 JAK1 KO (Supplemental Figure 4e, f), corresponding to the presence of large, greater than 4N cells with condensed mitotic chromatin visualized with DAPI staining by fluorescence microscopy (Supplemental Figure 4g). These mitotic tetraploid cells were 2.6-fold more abundant than typical 4N mitotic cells and were arrested following radiation treatment (Supplemental Figure 4d, h). Thus, reduced chromatin condensation and the enhanced pS10-H3 signal in 8N cells suggests that the M phase progression defect may in part be resolved by mitotic slippage and progression to tetraploidy.

Cell cycle progression initiated from the G1 phase was also examined in Cal27 or Detroit562 isogenic cell line pairs using a double thymidine block followed by immediate exposure to 8Gy. Remarkably, after synchronization Cal27 and Detroit562 JAK1 KOs displayed an identical accumulation of a G2/M population, while the isogenic controls completely arrested in the G1 phase (Figure 5i-l). As thymidine itself causes replication stress and DNA damage^20^, these results indicate that even mild genotoxic stress is capable of initiating a G2 arrest in cells with loss of JAK1 function. Following thymidine release, controls displayed accumulation of G2/M cells at 8 hours and restoration of a G1 population at 24 hours indicating successful mitosis. JAK1 KO cells also progressed through the cell cycle with accumulation of G1 populations after passing through G2/M, but with a greater portion of cells in the G2/M or tetraploid state compared to controls (Figure 5i-l). Following radiation, a greater population of G2/M or G2/M and tetraploid cells was observed in JAK1 KOs demonstrating sequential loss of cell cycle progression through both G2 and M phases (Figure 5i-l).

### Radiation-induced mitotic death is prevented by G2 cell cycle arrest

Mitotic catastrophe is the primary mode of cell death following exposure to ionizing radiation. The discovery of a prolonged G2/M phase associated with reduced micronuclei formation thus implies that loss of JAK1 function prevents mitotic entry and mitotic death. To test this possibility, we directly monitored cell fate with live cell imaging of thymidine synchronized and released control or JAK1 KO Cal27 cells treated with or without radiation (Figure 6a; Supplemental Videos 1a, b). Time lapse imaging was initiated 8 hours after 4Gy and continued for an additional 10 hours with visualization and scoring of individual mitoses. JAK1 KOs entered mitosis at a significantly reduced rate (15% vs. 4%; p<.01; Fig. 6c), providing clear evidence for a G2 progression defect. Moreover, mitotic cell death was significantly reduced in JAK1 KO cells compared to controls (11% vs 2%; p<.001; Figure 6c).

**Figure 6:**
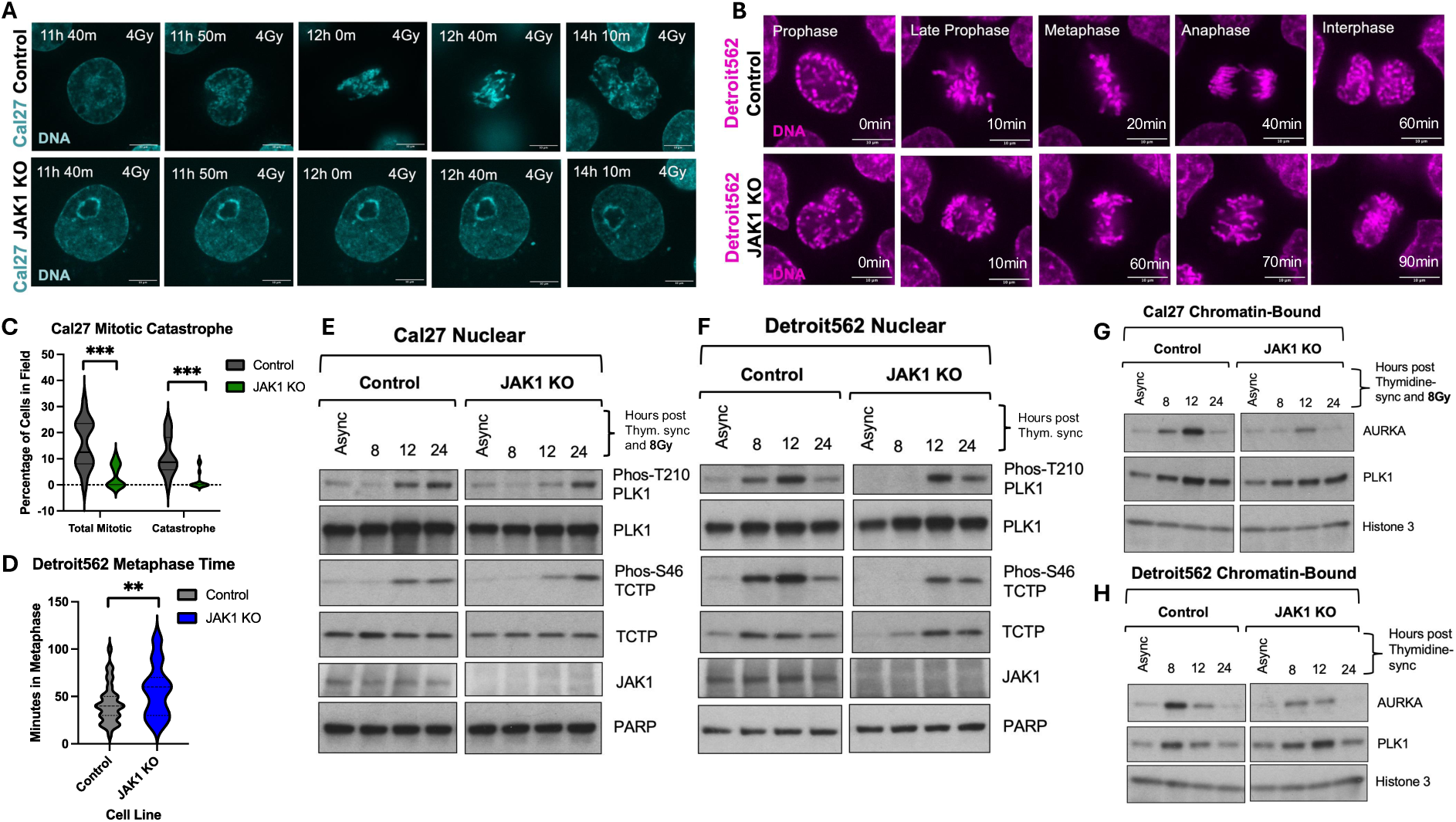
JAK1 dependent cell cycle progression and PLK1/Aurora Kinase A signaling. (**A**) Live cell imaging of synchronized and released Cal27 control of JAK1 KO cells following irradiation with 4Gy and visualized with NucSpot Red live cell dye for 10 hours. Representative time lapse images illustrate differences in progression to mitosis and mitotic death. Time stamp shown. (**B**) Live cell imaging for non-irradiated Detroit562 control or JAK1 KO cells illustrating delayed mitotic progression. Time from initiation of chromatin condensation is shown. (**C**) Quantification of mitotic entry and mitotic death corresponding to panel A. (**D**) Quantification of time spent in metaphase corresponding to panel B. (**E**, **F**) Western blots depicting phosphorylation changes of nuclear PLK1 and TCTP following synchronization and/or radiation treatment (8 Gy). (**G**,**H**) Western blots examining the presence of AURKA and PLK1 in the chromatin-bound fraction following synchronization and/or radiation treatment (8 Gy). ** indicates p<0.01. *** indicates p<0.005.

In the Detroit562 JAK1 KOs, the elevated baseline population of G2 cells provided the ability to use live cell imaging to monitor mitosis in the absence of radiation treatment (Figure 6b; Supplemental Videos 2a, b). Compared to controls, JAK1 KO cells frequently stalled in mitosis and had a prolonged metaphase that was on average 34% longer than controls (Figure 6d). In addition, failure of chromosome segregation in anaphase followed by collapse into a single cell was regularly observed (Figure 6b). In summary, live cell imaging experiments revealed that the G2 and metaphase progression defects caused by JAK1 KO reduce mitotic catastrophe and cell death.

### JAK1 regulates PLK1 and AURKA activity to enhance survival

To examine the signaling mechanisms that underlie enhanced G2 and M arrests in JAK1 KO cells, cell cycle protein activation states were determined over a 24-hour time course following thymidine synchronization and treatment with 8Gy. In the canonical pathway for DNA damage-induced G2 arrest, mitosis is prevented by phosphorylation and de-phosphorylation of CDK1. However, no differences in the T161 activating phosphorylation or the Y15 inhibitory phosphorylation were observed between control and JAK1 KO cells (Supplemental Figure 5a-d). Although a small increase in Cyclin B1 occurred with JAK1 KO, we interpret this finding to reflect the increased G2 population. In addition, no changes in activation of DNA-damage response proteins were observed (Supplemental Figure 5e-f). Together, these data suggest that hyper-activation of G2 and M arrests caused by loss of JAK function and independent of CDK1 signaling.

Polo-like kinase 1 (PLK1) is a serine-threonine kinase that participates in mitosis initiation through multiple effectors, resulting in centrosome maturation, bipolar mitotic spindle assembly, and activation of the anaphase-promoting complex^21^. We therefore examined PLK1 phosphorylation states using the nuclear fraction of thymidine synchronized Cal27 and Detroit562 control and JAK1 KO cells after treatment with 8Gy. In both, loss of JAK1 function delayed activation of PLK1 following thymidine release (Figure 6e, f) consistent with altered cell cycle kinetics (Figure 5j). While PLK1 activation occurred in control cells by 8-12 hours, full activation of PLK1 in JAK1 KO cells was not observed until 12-24 hours following treatment. This data is also consistent with the pS10 H3 data which shows reactivation of mitosis in JAK1 KO cells beginning at 24 hours (Figure 5d). Reduced phosphorylation of serine 46 of TCTP, a downstream target of PLK1^22^, was also observed and confirmed that loss of JAK1 function causes delayed activation of PLK1 (Figure 6f).

PLK1 is a signaling partner of Aurora Kinase A (AURKA), which has multiple functions at mitotic chromatin including phosphorylation of pS10-H3^23^ and regulation of sister chromatid cohesion^24^. Because we observed both PLK1 signaling changes and mitotic abnormalities, we hypothesized that AURKA chromatin localization would also be disrupted. Indeed, following thymidine release, chromatin bound AURKA, but not PLK1, was reduced in both the Cal27 (Figure 6g) and Detroit562 (Figure 6h) JAK1 KO cells compared to controls (Figure 6g-h). This loss of chromatin-localized AURKA during cell cycle progression is consistent with disruption of PLK1/AURKA signaling as a mechanism for G2 and M phase dysfunction in the setting of JAK1 KO.

The G2 cell cycle arrest induced by DNA double strand breaks enables recruitment of Rad51 to initiate homologous recombination (HR) dependent DNA repair. Notably, a 50% increase in Rad51 foci was observed at 4- and 8-hours after treatment with 4Gy in both Detroit562 and Cal27 JAK1 KOs (Figure 7a-d). This result indicates that the prolonged G2 arrest promotes extended access to the HR DNA repair pathway to improve cell survival. Bim expression, a marker and effector of cell death, was also reduced in synchronized JAK1 KO cells after radiation treatment (Figure 7e, f). Together, these results offer a more complete understanding of the downstream effects of JAK1 KO: reduced PLK1 and AURKA signaling, enhanced G2 arrest, metaphase progression defects, enhanced usage of HR, reduced mitotic death and micronuclei formation, and HNSCC radioresistance (Figure 7g).

**Figure 7:**
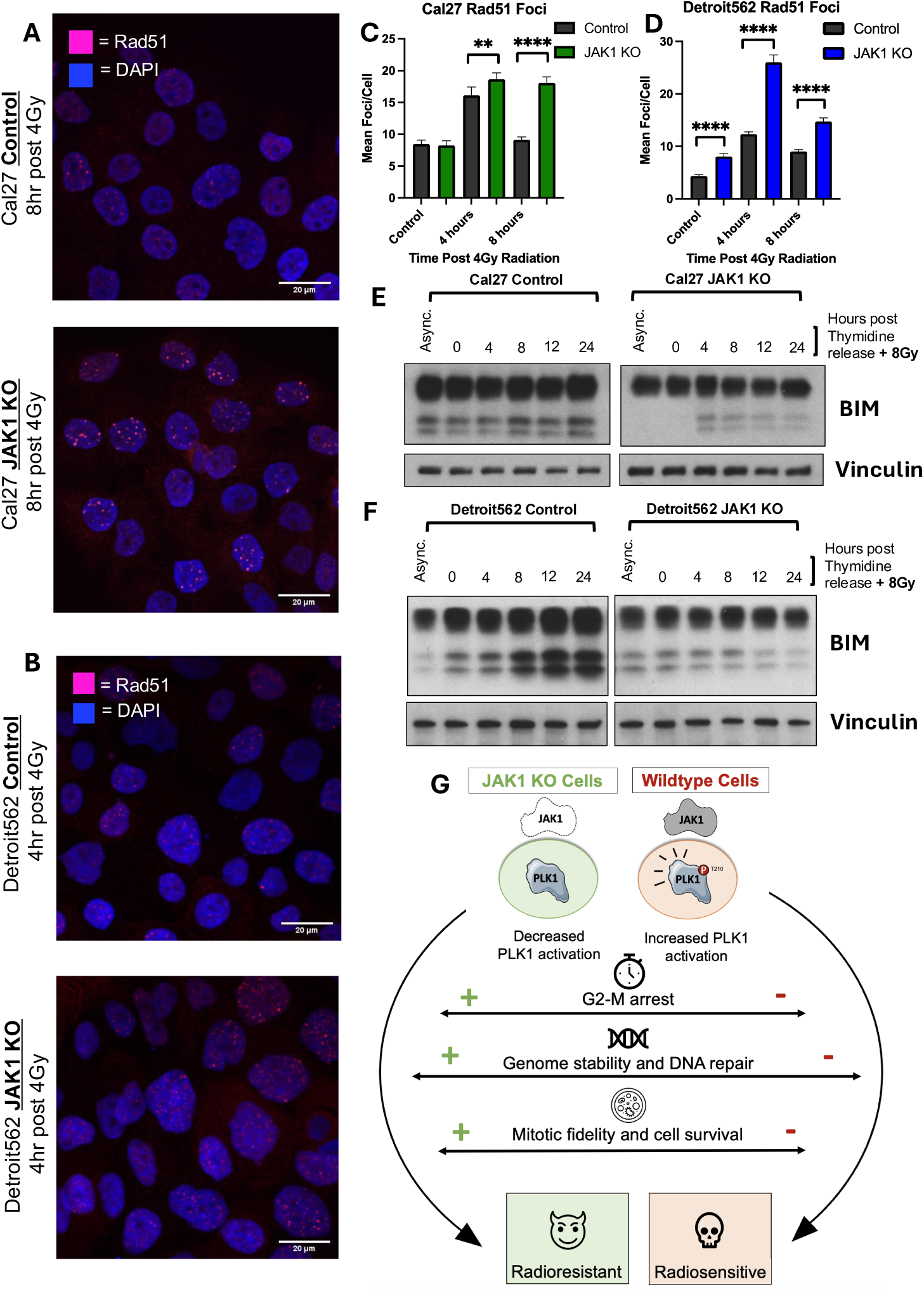
Differential induction of DNA repair and cell death markers with JAK1 KO. Representative images of Rad51 foci in (**A**) Cal27 or (**B**) Detroit562 control and JAK1 KOs at 8 hours following control or radiation (4Gy) treatment, respectively. (**C**,**D**) Quantification of mean foci per cell normalized for nuclear area for 4 and 8 hour time points after radiation. Time course of Bim protein expression in (**E**) Cal27 and (**F**) Detroit562 control and JAK1 KOs by Western blot after release from thymidine synchronization and radiation (8Gy) treatment. (**G**) Model of the cellular effects caused by JAK1 KO and that lead to resistance to radiation. ** indicates p<0.01. **** indicates p<0.001.

### Exploiting mitotic defects to overcome radioresistance

The radioresistance caused by JAK1 KO provides a new model for testing therapeutic strategies in tumors with altered cell cycle kinetics. We therefore investigated the effects of targeting Wee1, an established strategy to inhibit DNA damage-induced phosphorylation of CDK1^25,26^ and to bypass the protective G2/M arrest. While treatment with 500nM of the Wee1 inhibitor adavosertib decreased the radiation-induced G2 arrest of controls by 44.3% and 31.0% in Cal27 and Detroit562, respectively (Figure 8a, c), adavosertib did not abrogate the G2/M arrest in JAK1 KOs (Figure 8a-d; Supplemental Figure 6a-d). Inhibition of CDK1 phosphorylation by adavosertib was confirmed by western blotting (Supplemental Figure 6e), excluding redundant MYT1-dependent CDK1 phosphorylation. These results thus provide a mechanistic understanding for potential limitations of therapeutic strategies to activate CDK1, and underscore the value of identifying and modeling specific cell cycle progression defects.

**Figure 8:**
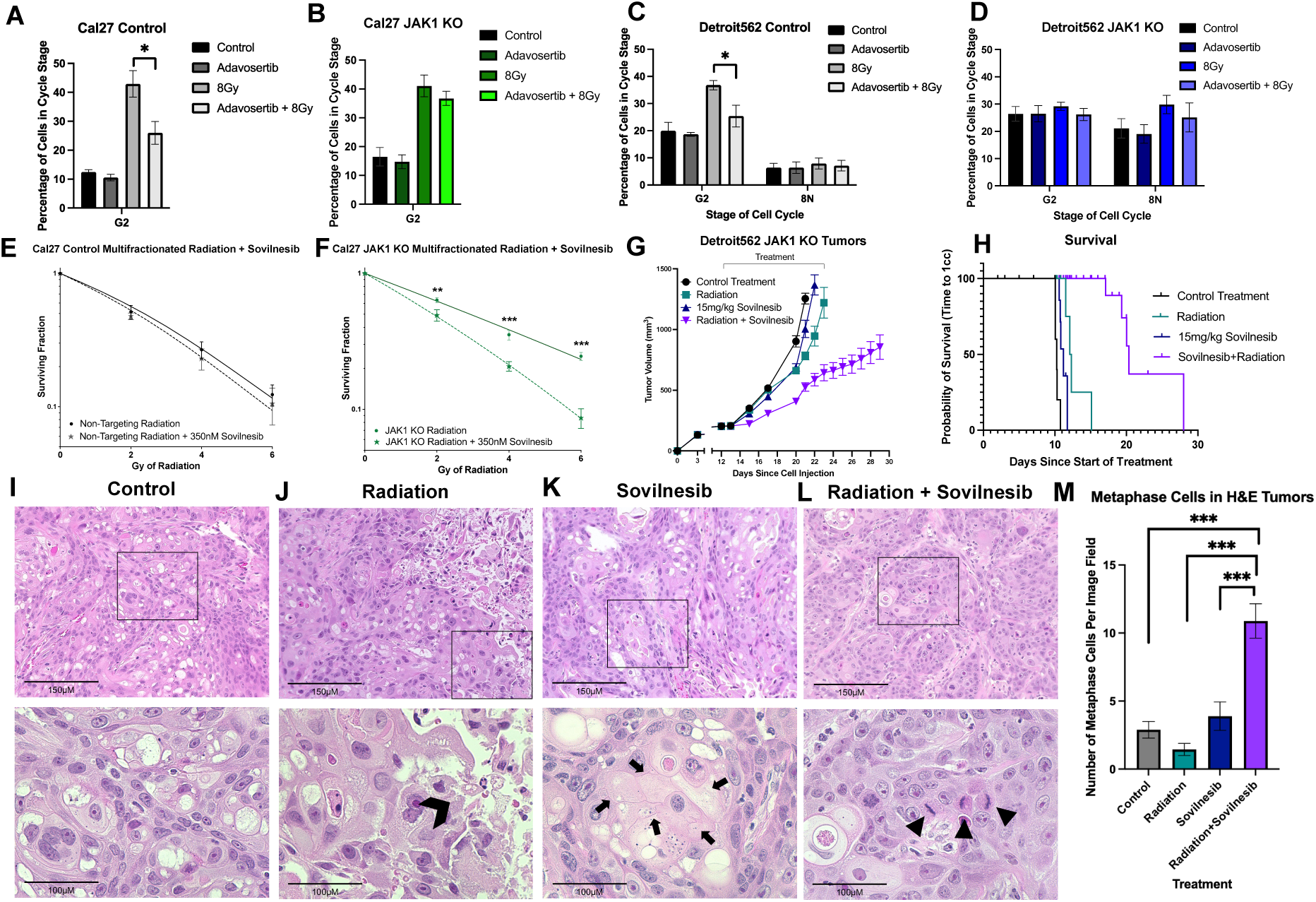
Mitotic progression defects confer sensitivity to Kif18A inhibition. Cell cycle distributions of (**A**,**B**) Cal27 or (**C**,**D**) Detroit562 control and JAK1 KO cells 8 hours following treatment with 500nM of adavosertib, 8Gy, or a combination of the two. Clonogenic survival of Cal27 (**E**) control or (**F**) JAK1 KOs treated with or without 350nM of sovilnesib and daily 2Gy fractions of radiation to achieve indicated doses is shown. (**G**) Tumor growth delay of Detroit562 JAK1 KO tumors in athymic nude mice treated with either vehicle, daily radiation (2Gy x 9), solvinesib (15mg/kg), or solvinesib plus radiation. (**H**) Kaplan Meier survival curves illustrating time to tumor growth of 1cm^3^ (and sacrifice). Representative haematoxylin and eosin (H&E) staining of (**I**) control, (**J**) radiation, (**K**) sovilnesib, and (**L**) radiation plus sovilnesib groups. Region of interest in 20X images (top panels) are indicated by black squares and presented at 60X magnification. The black chevron indicates radiation cell death, black arrows identify regions of cell death with loss of nuclei, and back arrow show cells arrested in metaphase. (**M**) The number of metaphase cells in each 60x image were quantified for 3 images per each of the 3 tumors per experimental group. * indicates p< 0.05. ** indicates p<0.01. *** indicates p<0.005.

Kif18a is a kinesin ATPase that activates microtubule depolymerization to shorten kinetochores and maintain kinetochore tension. Remarkably, a dependency on the function of Kif18a has been demonstrated for cells with whole genome duplication or aneuploidy^27,28^. Given the metaphase progression defect and tendency for JAK1 KO cells to progress to tetraploidy, we hypothesized that Kif18a inhibition would enhance the effect of radiation. The small molecule Kif18a inhibitor sovilnesib (350nM) was therefore first tested *in vitro* using a multi-fractionated radiation treatment of 2Gy/day for three days in Cal27 cells. In controls, the combination of sovilnesib and radiation had only a small effect on radiosensitivity (Figure 8e). In contrast, significant radiosensitization of JAK1 KOs was observed (Figure 8f). The effect of sovilnesib was also tested in mice bearing Detroit562 JAK1 KO tumors and treated with daily radiation, 15mg/kg of sovilnesib, or combined treatment. JAK1 KO tumors again showed no significant growth delay with radiation, and treatment with sovilnesib alone also showed no effect on growth. However, delivery of sovilnesib concurrently with radiation was synergistic, resulting in an average 11-day delay in tumor growth (Figure 8g-h). Treatment with radiation and sovilnesib was well tolerated and no weight loss was observed (Supplemental Figure 7a). Together these results indicate that addition of sovilnesib to radiation is an effective strategy to radiosensitize tumor cells with loss of JAK1 function. In a broader perspective, they also suggest that cell cycle progression defects that enable cell survival but cause mitotic stress can be countered by pharmacologic inhibition of Kif18a.

To further characterize tumor responses, we performed haematoxylin and eosin (H&E) staining on three Detroit562 JAK1 KO tumors from each treatment group (Figure 8i-l, Supplemental Figure 7d-g). Control tumors showed high cellularity and disorganized tissue architecture, consistent with a highly aggressive tumor (Figure 8i; Supplemental Figure 7d). In the radiation monotherapy group, tumors displayed pockets of cell debris consistent with radiation-induced tumor necrosis (Figure 8j; Supplemental Figure 7e). Sovilnesib monotherapy treated tumors were also less cellular than controls with pockets of eosin positive cells lacking a cell nucleus (Figure 8k; Supplemental Figure 7f). In the combined therapy group, an accumulation of cells undergoing mitosis was observed alongside evidence of coagulative necrosis (Figure 8l; Supplemental Figure 7g). In particular, there was a significant increase in mitotic metaphase arrested cells in the combined therapy group compared to all other treatment groups (p<.01; Figure 8m), consistent with inhibition of the metaphase to anaphase transition^29^. Together, these results provide compelling evidence that although cell cycle progression defects confer resistance to DNA damage, Kif18a inhibition exploits a mechanistically linked and inherent M phase vulnerability to enhance the efficacy of radiation therapy.

## DISCUSSION

Pooled gRNA CRISPR-Cas9 screening in two HNSCC cell types revealed that loss of JAK1 function promotes radioresistance. This finding was validated in both *in vitro* clonogenic and *in vivo* xenograft models using isogenic control and JAK1 KO Cal27 and Detroit562 cell lines. We find that JAK1 KO causes an unexpected enhancement of the G2 cell cycle arrest following exposure to radiation. This cell cycle effect was confirmed with multiple techniques in asynchronized, thymidine synchronized, and Edu labeled cells. Loss of JAK1 also induces defects in mitotic progression, specifically a protracted metaphase time, and both cell cycle deficits are associated with attenuated activation of PLK1. In the setting of DNA damage, the enhanced G2/M arrest confers greater access to HR for DNA repair, a decrease in mitotic catastrophe and micronuclei formation, and a decrease in Bim signaling. Although we found that CDK1 activation through Wee1 inhibition was not able to reverse the G2/M arrest in JAK1 KOs, we demonstrate that inhibition of the mitotic kinesin, Kif18a, exploits an associated mitotic progression defect and restores sensitivity to radiation.

Screening with the kinome-focused CRISPR-Cas9 gRNA library revealed both known and unknown regulators of the cellular response to radiation. We found that gRNAs targeting *ATM* or *PRKDC* (coding for DNA-PK) were depleted in both screens, indicating that loss of function confers sensitivity to radiation. This is consistent with the known roles of ATM and DNA-PK in the DNA damage response. Indeed, loss of ATM has been shown to broadly confer sensitivity to DNA damaging agents including cisplatin, radiation, and PARP inhibitors^30^. The identification of *PRKDC* likewise aligns with experimental findings that pharmacologic inhibition of DNA-PK sensitizes multiple HNSCC cells to radiation^12^. The screening results also suggest that loss of NFKB signaling genes causes radioresistance. These results agree with prior studies showing that activation of NFKB by Toll-like receptors enhances sensitivity to fractionated radiation^13,14^. Remarkably, we also found JAK1 to be the top hit promoting radioresistance in both Cal27 and Detroit562 cells. This finding led us to the discovery of an intrinsic role for JAK1 in the cellular response to radiation. Although loss of JAK1 has been shown to cause resistance to immunotherapy^31^, the work presented herein describes a previously unrecognized mechanism by which loss of JAK1 function causes therapeutic resistance to a second fundamental oncology modality: agents that cause DNA damage. Together the CRISPR screening results demonstrate success of this approach for identifying both established and previously unrecognized pathways that regulate the tumor cell response to radiation.

The role of G2 cell cycle arrest in promoting DNA repair and enhancing radioresistance has been understood for more than 40 years^32^. Activation of G2 cell cycle progression, especially in p53 mutant tumor cells, has therefore been pursued as a therapeutic strategy to enhance the effects of DNA damage^33,34^. This approach is well illustrated by inhibition of Wee1, a kinase that suppresses CDK1 activity, and can be targeted to promote G2 progression and cell death after DNA damage^25,34^. Our results indicate that beyond this canonical regulatory pathway, a second JAK1-dependent pathway is also operational in the setting of DNA damage. We report that JAK1 KO enhances G2 arrest by delaying activation of PLK1 and disrupting AURKA trafficking to chromatin, consistent with the roles for these proteins in overcoming the DNA damage-induced G2 arrest^35^. In addition, the JAK1 KO phenotype is similar to that for reduced AURKA activity, with formation of giant polyploid and therapeutically resistant tumor cells^36^. Although neither PLK1 nor AURKA were identified as hits in the genetic screen, this is likely due to their essential nature. In contrast, JAK1 is not essential, with KO only impacting the dynamic function of downstream proteins and enabling prolongation of G2 without eliminating essential mitotic functions. Because mass spectrometry studies have identified 35 PLK1 phosphorylation sites, including tyrosine sites^37^, additional work to elucidate whether JAK1 to PLK1 signaling is direct or indirect will be required. Although JAK1 has not previously been reported to affect PLK1 activation, JAK2 has recently been found to regulate CHK2-dependent PLK1 activation and promote timely mitosis in U2OS and HeLa cells^38^. Our results in HNSCC, however, did not demonstrate an effect of JAK1 KO on ATM or ATR (upstream regulators of CHK1 and CHK2; Supplemental Figure 5). Furthermore, ATR, JAK2, CHK1, and CHK2 were not identified as potential targets by the genetic screens. Collectively, this data raises the possibility that JAKs may have conserved and complementary mechanistic roles in regulating the cell cycle in the context of DNA damage.

Suppression of the radiation-induced G2/M arrest using Wee1 inhibitors is a therapeutic strategy actively being evaluated in clinical trials^26,39^. Surprisingly we found that, unlike controls, Wee1-dependent CDK1 activation is not capable of driving G2 progression following DNA damage in the setting of JAK1 KO. These results confirm that JAK1’s role is independent of the canonical CDK1-dependent G2 checkpoint, and further highlight the crucial role of PLK1 activity for commitment to mitosis^21^. This interpretation is also supported by the timeline for mitotic chromatin formation after radiation, as indicated by pS10-H3 (Fig. 5c-d, g-h), which coincides precisely with reactivation of PLK1 phosphorylation. Given that JAK1 loss of function mutations are well known to emerge after anti-PD1 therapy in melanoma^33^, and have also been reported in HNSCC^40^, NSCLC^41^, and pre-clinical models of colon cancer^42^, it is likely that tumors frequently harbor tumor subpopulations with JAK1 inactivation. In addition, while JAK1 mutation *per se* is detectable through exome sequencing, other pathway mutations or epigenetic changes that impact JAK1 function, such as suppression of JAK1 activity by the PTPN2 phosphatase^31^, represent less readily identifiable mechanisms. Tumor heterogeneity with rare populations of tumor cells with JAK1 inactivation thus represent not only a putative mechanism for therapeutic resistance to Wee1 inhibition but also a second, dominant pathway for regulating cell cycle progression and enabling resistance to radiation therapy and other DNA damaging agents.

A prominent observation from our studies is that the prolonged G2/M arrest caused by JAK1 KO results in reduced formation of micronuclei, presumably reducing cell death caused by stochastic inheritance of micronuclei by daughter cells^43^. Previous work has shown that micronuclei formation following chromosome mis-segregation events predispose cells to chromothripsis and amplify radiation-induced genome instability^44^, and reduced micronuclei formation also limits cytosolic DNA damage associated molecular pattern detection by cGAS-STING signaling, a second pathway that modifies tumor cell radioresistivity^3,44,45^. Thus, reduction of micronuclei formation has several downstream consequences for tumor cell survival. Herein we demonstrate that the prolonged G2/M arrest also reduces protein levels of Bim, a BH3 only and pro-death Bcl2 family member. While Bim function is most often associated with apoptosis, its phosphorylation by AURKA during mitosis^46^ and its requirement for cell death induced my microtubule targeting agents^47^ suggests that reduced Bim signaling is another mechanism that limits radiation-induced mitotic catastrophe.

A phenotypically associated consequence of reduced PLK1 and AURKA activity is a defect in mitotic progression. AURKA phosphorylates mitotic spindle proteins and is critical for spindle formation and stability, while PLK1 activity ensures that kinetochore-microtubule attachments are correctly formed and stable^48^. We hypothesized that reduced PLK1/AURKA signaling would therefore induce mitotic stress and a mitotic arrest after DNA damage, and in agreement, observed the emergence of tetraploid cell populations by flow cytometry and fewer mitoses with live cell imaging in JAK1 KOs following radiation exposure. Based on this new insight, we advanced Kif18a inhibitors to test in combination with radiation. Kif18a is a non-essential kinesin ATPase that regulates the length of kinetochores, maintains kinetochore tension to satisfy the spindle assembly checkpoint, and has been shown to be required for division of aneuploid tumor cells^42,43^. Our results demonstrate that inhibition of Kif18a does indeed enhance radiosensitivity of HNSCC with JAK1 KO, but has only a small effect on isogenic controls. In xenograft tumor experiments, an elevated number of metaphase cells was observed following radiation and sovilnesib treatment, consistent with the anticipated Kif18a inhibitor mechanism of action. As Kif18a inhibitors have recently advanced to phase I clinical trials (NCT06084416), targeting Kif18a in combination with DNA damaging agents thus provides an alternative, mechanism-based approach to neutralize the enhanced survival conferred by DNA damage-induced G2 arrest.

The discovery that *loss* of JAK1 function is a major driver of tumor cell-intrinsic resistance to ionizing radiation ostensibly contrasts with studies implicating that upregulation of IFN-JAK-STAT signaling promotes resistance to radiation therapy. For example, seminal work from Minn, Weichselbaum, and colleagues has shown that radiation and chemotherapy resistant tumors upregulate interferon-related gene expression^49^, and that resistance to radiation is acquired by overexpression of STAT1^50^. However, short term STAT1 activation by radiation or interferon has also been suggested to cause cell death^51^, and other studies propose that interferon beta is essential to amplify the DNA damage response and promote senescence^52^. These seemingly contradictory results can be reconciled when considering the timing and duration of the interferon response. Our data is consistent with an initial phase of interferon signaling causing cytotoxicity and cell death, whereas long term interferon or inflammatory signaling leads to cellular adaptation and therapeutic resistance. In this framework, both acute loss of JAK1 and adaptation to chronic inflammatory signaling mediate the same effect: an inability for the cell to initiate cell death programs following exposure to radiation. Intriguingly, a similar paradox exists with resistance to immunotherapy, where signatures of prolonged interferon signaling and STAT1 activation are markers of immune dysfunction and found in cells resistant to immune checkpoint blockade (ICB)^53,54^. A recent clinical trial has explored this phenomenon using JAK inhibitors in combination with ICB with the goal of relieving chronic inflammatory signaling to re-sensitize cells to cytotoxic T cell signaling with promising results^55^. JAK1 inhibition to reduce tumor proliferation has also been considered as an experimental intervention for solid tumors in the absence of immunotherapy^56,57^, and FDA approved JAK1 inhibitors are frequently used for the treatment of inflammatory disease. However, given our data demonstrating a tumor cell survival benefit with loss of JAK1 function, our work raises questions regarding deployment of this inhibitor class in combination with DNA damaging agents in oncology settings, and suggests that clinical studies of this combination should proceed with caution.

In summary, we report the results of pooled genetic CRISPR-Cas9 screening campaign designed to identify signaling pathways that alter cellular survival following exposure to ionizing radiation. The unexpected discovery that JAK1 impacts cell cycle progression offers a new perspective regarding control of a fundamental cellular mechanism long understood to cause therapeutic resistance to DNA damage. Subsequent exploration of the underlying cell biology in JAK1 deficient cells uncovered a mitotic vulnerability that can be targeted through Kif18a inhibition, thus offering new strategies for therapeutic intervention in oncology.

## MATERIALS AND METHODS

### Cell Lines and treatments

Cal27 (CVCL_1107) and Detroit562 (CVCL_1171) head and neck squamous cell carcinoma cell lines were obtained from the American Type Culture Collection (ATCC). HEK393T (CVCL_0063) cells used for lentivirus generation were a gift from Dr. Ryan Jensen (Yale University, New Haven, CT). Cell lines were maintained in DMEM media (Gibco Cat# 11965092) supplemented with 10% FBS (Gibco Cat# 16000-044). All cells were kept in a humidified incubator with 5% CO_2_ and were kept in culture for no longer than 10 passages from defrosting. Cells were routinely tested for mycoplasma contamination using the MycoAlert Mycoplasma Detection Kit (Lonza Cat# LT07-318). HNSCC cell lines were validated by STR profiling by ATCC. Radiation was delivered locally using a X-RAD 320kV orthovoltage unit at a dose rate of 2.3 Gy/min with a 2mm aluminium filter (PXi, Precision XRay). Quality assurance was performed monthly using a P.T.W. 0.3 cm^3^ ionization chamber calibrated to NIST standards and quarterly dosimetry using thermoluminescent dosimeter-based or ferrous sulphate-based dosimeters. Abrocitinib (HY-107429), Sovilnesib (HY-132840) and Thymidine (HY-N1150) were purchased from MedChem Express (Monmouth Junction, NJ USA). Adavosertib was a gift from Dr. Barbara Burtness (Yale University, New Haven, CT). The concentration of drugs used in this study can be found in Supplemental Methods Table 1. Dose and treatment regimens for the drugs are described in the figure legends.

### Lentivirus production

Virus production for Cas9 plasmid, JAK1 KO gRNA plasmid and the pooled library was achieved as previously described^58^.

### Cas9 and JAK1 knockout cell line generation

Cal27 and Detroit562 cells were transduced with lentivirus containing a plasmid coding for Cas9 (lentiCas9-Blast; Addgene Cat#52962) as previously described^59^. Lentivirus was produced as above, and cells were transduced followed by Blasticidin S (Gibco Cat# A11139-03) selection. Single clones were then grown from the polyclonal pool and used for subsequent knockout or screening. gRNAs targeting JAK1 were cloned into the lentiGuide-Puro plasmid (Addgene #52963) and introduced into the cell via lentiviral transduction. Lentivirus was produced as above, and cells were transduced followed by Puromycin (Gibco Cat# A11138-03) selection. gRNA sequences are listed in Supplemental Table 2.

### CRISPR Screen

The Human Kinome CRISPR Knockout Library (Brunello) was purchased from Addgene (# 1000000082) and was generated as previously described^10^. Library amplification and sequencing were performed according to the manufacturer’s protocol (Addgene/Broad Institute). Prior to performing the screen, the human Kinome library representation was validated (Suppl. Figure 1). For each replicate of the pooled screen ∼10 x 10^6^ Cas9-expressing cells were transduced at a MOI of ∼0.3 with lentivirus produced as described above using from the aforementioned CRISPR-KO (#75312 (gRNAs 1-4) and #75313 (gRNAs 5-8)) library and selected with 2 µg/mL puromycin (Gibco) for 7 days. After puromycin selection, pooled cells were irradiated (2 Gy daily for 4 consecutive days) with a Precision X-Ray 320kV orthovoltage unit. Fourteen days after the final radiation treatment, cells were collected with unirradiated cells from the same experiment used as a control. Genomic DNA isolation was performed using QIAamp DNA Blood Midi kit (Cat# 51104. Qiagen, Hilden, Germany) and sgRNA sequence were amplified using a PCR reaction according to the manufacturer’s protocol (Addgene/Broad Institute) with maintenance of ∼400x coverage of the Human Kinome library. All PCR reactions were performed using Ex Taq DNA polymerase (Clontech, RR001A). Primers and barcode sequence are listed in the Tables 2-3. Sequencing was performed with NovaSeq paired-end 100bp reads. Reads were aligned to index sequences using the Bowtie aligner, and a maximum of one mismatch was allowed in the 20bp sgRNA sequence. The number of uniquely aligned reads for each library sequence was calculated after alignment for each of the three biologically independent replicates. Differential sgRNA expression was analyzed in R using the Model-based Analysis of Genome-wide CRISPR/Cas9 Knockout method ^11^. FDR-corrected p-value < 0.05 was considered statistically significant.

### Western blots

Primary Antibodies are listed in Supplemental Methods Table 4. Cell samples were lysed on ice in buffer containing 1% TritonX-100, 10mM of EDTA (pH 8.0), 25mM Tris (pH 7.4), 15% glycerol, 1% Phosphatase Inhibitor Cocktail II (Sigma Cat# P5726-5ML), 1% Phosphatase Inhibitor Cocktail III (Sigma Cat# P0044-5ML), and supplemented with one Complete Mini Protease Inhibitor (Roche Cat# 11836170001). Samples were run on a 4-20% polyacrylamide gel and transferred to a nitrocellulose membrane via a semi-dry transfer. Membranes were blocked with 5% skim milk and then incubated with primary antibodies at listed concentrations (Supplemental Methods Table 4) overnight at 4°C. Nitrocellulose-bound primary antibodies were then detected with anti-rabbit IgG horseradish peroxidase-linked antibody or anti-mouse IgG-horseradish peroxidase-linked antibody (EMD-Millipore) and detected by Amersham ECL detection reagent (GE Healthcare).

### Xenograft tumors studies

For *in vivo* studies shown in Figure 2, five million Detroit562 control or JAK1 KO cells were injected into the flanks of 4-6-week-old athymic Swiss nu/nu mice (Envigo) to initiate tumors. When tumors reached 200mm^3^, mice for each tumor group were randomized to receive either sham irradiation or a total of 20 Gy administered in daily 2 Gy fractions with a 2-day break after administration of the first 10 Gy. Radiation was administered using a Precision X-ray 320-kV orthovoltage unit. For *in vivo* studies shown in Figure 8, Detroit562 JAK1 KO tumors were initiated as described above. When tumors reached 200mm^3^, mice were treated with a multifractionated radiation regimen of 2Gy per day, 15mg/kg of sovilnesib, a combined treatment of radiation and sovilnesib, or a sham regimen. Because tumors in both monotherapy groups grew through the planned two weeks of monotherapy, sovilnesib monotherapy mice were sacrificed after 12 doses of sovilnesib, radiation monotherapy mice were sacrificed after 9 doses of 2Gy (18Gy total), and combination treatment mice received a total of 12 doses of sovilnesib and multifractionated 18Gy. Tumor size was measured, and volume was calculated according to the formula π/6 x (length) x (width)^2^. Mice were sacrificed when tumors reached 1000mm^3^. Tumor growth delay was calculated as time to reach 500mm^3^ as previously described^3^. Data are expressed as the mean volume ± SEM tumor volume. Group sizes are specified in the respective figures.

### Clonogenic assays

For single-fraction radiation clonogenics, 1000-2000 cells were seeded in 6 well plates and irradiated at indicated doses. For multifractionated radiation clonogenics, 100,000 cells were seeded in 60 mm dishes, irradiated as described, and then re-seeded at a density of 1000-2000 cells per well in a 6 well plate. Seeding for sovilnesib-treated clonogenics can be found in Supplementary Methods. All clonogenics were incubated for 14 days after seeding and then stained 0.25% crystal violet and 80% methanol. Colonies with greater than 50 cells were hand counted. The surviving fraction of each sample was calculated as the ratio between the number of colonies counted divided by number of cells seeded and the plating efficiency, thus normalizing for plating efficiency differences with treatment. Clonogenic survival differences for each treatment were compared using survival curves generated from the linear quadratic equation as previously described.

### Flow cytometry studies

Cell cycle distribution analysis, phos-Ser10 H3 mitosis assays, EdU assays and double thymidine synchronization were performed according to manufacturers’ protocols. Please see supplementary materials for further details.

### Micronuclei formation assay

Twenty-thousand cells were seeded into each chamber of a Lab-TekII chamber slide system (Thermo Scientific Cat#154526), incubated for 48 hours, and irradiated with 4 Gy. For experiments utilizing Abrocitinib (MedChem Express Cat# HY-107429), cells were incubated with indicated doses of Abrocitinib for one hour prior to radiation treatment. Inhibitor was left in the wells during subsequent incubation.

Twenty-four hours after radiation treatment, cells were fixed using 4% Paraformaldehyde in PBS and then permeabilized in 1% PBS-triton for 30 minutes at room temperature. Samples were incubated in blocking buffer containing 10% horse serum, 10% goat serum (Cat# G9023 and Cat# H0146, respectively. Sigma-Aldrich, St. Louis, MO USA) and eighty percent 1% PBS-triton for one hour at room temperature. Samples were washed once with 1% PBS-triton and then incubated in Anti-cGAS primary antibody overnight at 4°C. The next morning, samples were washed 3 times with 1% PBS-triton and then incubated in Alexa Flour 488 goat anti-rabbit antibody at room temperature for one hour (Supplemental Methods Table 4). Samples were washed 3 times with 1% PBS-triton and then mounted in ProLong Gold Antifade Mountant containing DAPI (Invitrogen Cat# P36935). Cells were imaged on an EVOS M5000 Fluorescent Microscope (AMF5000. Thermo Fisher Scientific Inc., Waltham, MA USA) with 60x objective. Total cell nuclei, micronuclei, and cGAS-positive micronuclei were manually counted. Raw numbers of cell nuclei, micronuclei, and cGAS-positive micronuclei were used to determine the ratio of micronuclei: nuclei per image.

### Rad51 foci formation assay

One hundred and ten thousand cells were seeded on glass coverslips in the bottom of 6 well plates and incubated for 48 hours. Cells were irradiated with 4Gy and fixed at the indicated timepoints in PTS solution for 15 minutes at room temperature. PTS solution contains 3 parts 4% PFA in PBS and one part 2% Triton + 32% Sucrose in PBS. Samples were rinsed once in PBS and then incubated in a 1:1 solution of ice cold Methanol: Acetone at 4^0^C for 10 minutes. Samples were rinsed in PBS and then incubated in blocking buffer containing 10% goat serum, 10% horse serum, and eighty percent 0.5% PBS-triton overnight at 4^0^C. Samples were washed once with 0.5% PBS-triton and then incubated with 1:300 anti-Rad51 overnight at 4^0^C (Suppl. Methods Table 4). Cells were rinsed twice with 0.5% PBS-triton and then incubated in secondary antibody as described above. Cells were washed 3 times and then counter-stained with a mixture (1:250 dilution) of 4′,6-diamidino-2-phenylindole (DaPi. Cat# D9542. Sigma-Aldrich, St. Louis, MO USA) and bisbenzimide H 33342 trihydrochloride solution (Hoechst 33342. Cat# B2261. Sigma-Aldrich, St. Louis, MO USA) in 0.5% PBS-triton for 5 minutes. Samples were rinsed three times with 0.5% PBS-triton and then mounted in Dako Florescence Mounting Media (Agilent Cat#S3032). Samples were imaged on a TiE Inverted Nikon spinning disk confocal microscope with 60x objective. Rad51 foci were counted The Focinator v2 software as previously described^60^.

### RNA Sequencing

Cells were split at passage 2 into three T75 flasks, one for each biological replicate. Two million cells from each T75 flask were then plated into 10cm dishes on separate, subsequent days. Forty-eight hours after seeding, cells were washed with PBS, tyrpsinized, and counted for RNA isolation. RNA was isolated using the Qiagen RNeasy kit (Cat#74104) according to manufacturer’s instructions. Samples were sent to the Yale Center for Genome Analysis for poly-A RNA sequencing (HiSeq paired-end, 100bp) using NovaSeq.

### Live Cell Imaging

Eighty thousand Cal27 or Detroit562 control and JAK1 KO cells were seeded into each chamber of a glass bottomed Ibidi imaging system (Cat# 80287). Cells were subjected to thymidine synchronization as previously described. For Cal27 cells, cells were irradiated with 4 Gy immediately following thymidine release. Seven hours following thymidine release, cells were stained with NucSpot Live 650 Nuclear Stain (Biotium Cat# 40082) for one hour. Following staining, cells were imaged overnight on a Nikon Crest X-LightV3 spinning disk confocal microscope with a sCMOS camera (Teledyne Kinetix, 10 MP, 6.5 um pixel) with 10-minute imaging intervals at 37°C and 5% CO_2_. Dye was imaged using 637 nm excitation light from an AURA light engine (Lumencor), a Semrock FF421/194/567/659/776-DI01 dichroic, and a Chroma ET665lp filter. Analysis of live cell data was performed using FIJI software.

### Statistical analysis

Results are expressed as mean ± standard error (S.E.) unless otherwise indicated. The Microsoft Excel (version 16.36) was used for statistical data analysis. GraphPad v 10.1.0 (264) was used for the statistical data analysis to compare the survival curves (time to reach 500mm3; Fig. 2).

Statistically significant differences in between-group comparisons were defined at a significance level of P-value ≤ 0.05 in 2-tail student’s t-Test of two-samples of unequal variance (heteroscedastic) or in Log-rank (Mantel-Cox) test.

### Study approval

All experimental procedures for animals were approved in accordance with Institutional Animal Care and Use Committee and Yale university institutional guidelines for animal care and ethics and guidelines for the welfare and use of animals in cancer research.

## Supporting information

Cal27 Control 4Gy Live Cell

Cal27 JAK1KO 4Gy Live Cell

Detroit562 Control Live Cell

Detroit562 JAK1KO Live Cell

Cal27 RNA Sequencing

Detroit562 RNA Sequencing

## Acknowledgements

This work was funded by the Project 1 of the Yale SPORE in HN Cancer P50DE030707 (J.N.C.) and the Yale Cancer Center Team Challenge Award. We thank Yale Flow Cytometry for their assistance with and use of the Cytoflex flow cytometer and the Yale Irradiator Shared Resource, both supported in part by the NCI Cancer Center Support Grant (P30CA016359). We thank the Yale West Campus Imaging Core for the support and assistance in this work as well as the Yale Center for Genome Analysis Core for their help with sequencing and analysis (S10OD030363).

## Author Contributions

Conception and design: V.K., M.B., J.N.C.

Development of methodology: V.K., M.B., W.G., C.P., J.N.C.

Acquisition of data: V.K., M.B., W.G., N.A., H. L., C.P., J.N.C.

Analysis and interpretation of data: V.K., M.B., W.G., N.A., H. L., L.K., J.N.C.

Writing original draft: V.K., J.N.C.

Writing, review, and/or revision of the manuscript: V.K., M.B., W.G., N.A., H. L., C.P., J.N.C.

Study supervision: J.N.C.

## Data and Material Availability

All data are available in the main text or the supplementary materials.

