## Supplemental Data and Methods for "Loss of JAK1 Function Causes G2/M Cell Cycle Defects Vulnerable to Kif18a Inhibition"

The PDF file includes:

Fig. S1 to S7

Table S1 to S2

Supplementary material and methods with methods tables T1 to T4

Other Supplementary materials for this manuscript includes the following: Supplemental Videos 1 and 2 for live cell imaging.

### SUPPLEMENTAL FIGURES AND TABLES

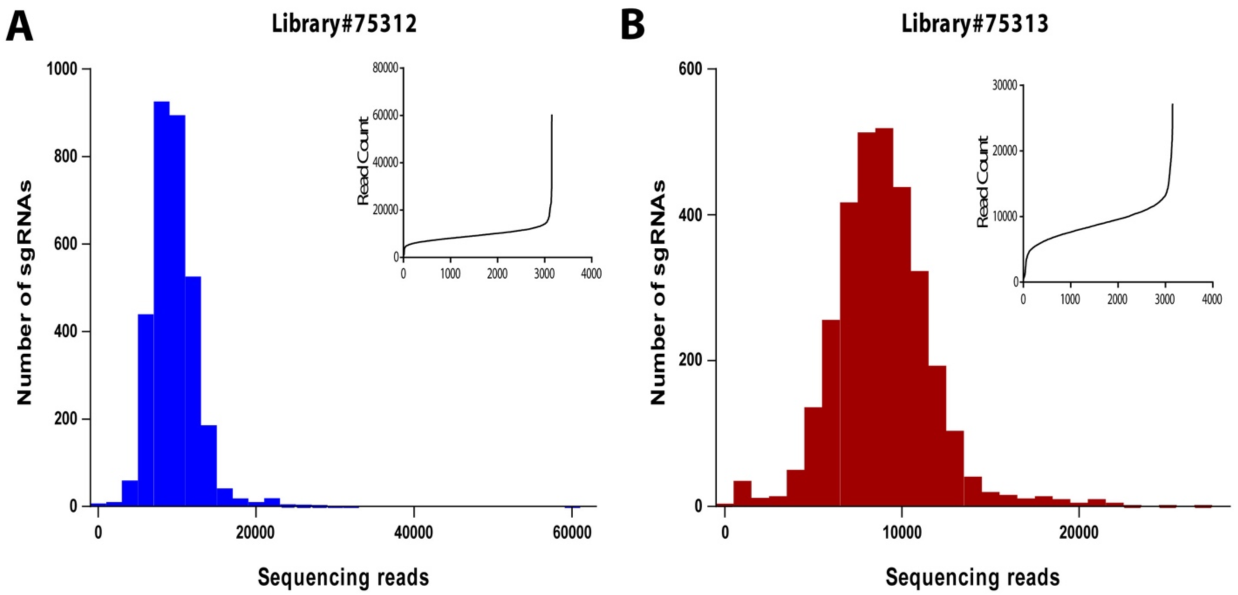


**Supplementary Figure 1. Sequencing of human Kinome CRISPR KO library cloned into lentiGuide-Puro.** Representation of sgRNA reads was verified with Next Generation Sequencing (NGS). Histograms of sgRNA representation of each half-library with accumulative distribution of sequencing reads. **(a)** Human Kinome CRISPR Knockout #75312 (gRNAs 1-4) and **(b)** Human Kinome CRISPR Knockout #75313 (gRNAs 5-8) in lentiGuide-Puro.

**Supplemental Table 1: Top 20 Kinome-Wide CRISPR Screen Hits in Cal27 and Detroit562 Cell Lines.** (A) Cal27 and (B) Detroit562 cell lines were transduced with a kinome-wide gRNA library and were split into two treated and untreated groups. The irradiated group was treated with 4 doses of 2Gy of radiation even 24 hours, and the unirradiated group was treated with a sham regimen. Two weeks after the final radiation dose, the cells were harvested, and amplified gRNAs were sent for deep sequencing to determine gRNAs that were enriched or depleted between the irradiated and unirradiated groups. The top 20 enriched and depleted gRNAs for the indicated genes are shown for both cell lines. FDR<0.01 was considered significant.

**A**

| CAL27 | | | | | |
|---|---|---|---|---|---|
| **id** | **neg\|p-value** | **neg\|fdr** | **id** | **pos\|p-value** | **pos\|fdr** |
| STK11 | 4.96E-06 | 0.000421 | JAK1 | 4.96E-06 | 0.000758 |
| ATM | 4.96E-06 | 0.000421 | ALDH18A1 | 4.96E-06 | 0.000758 |
| MAP2K7 | 4.96E-06 | 0.000421 | IRAK1 | 4.96E-06 | 0.000758 |
| PRKDC | 4.96E-06 | 0.000421 | TYK2 | 4.96E-06 | 0.000758 |
| HUS1 | 4.96E-06 | 0.000421 | IKBKB | 1.49E-05 | 0.001894 |
| SCYL1 | 4.96E-06 | 0.000421 | CHUK | 0.00021319 | 0.023268 |
| MAP3K7 | 4.96E-06 | 0.000421 | CLK1 | 0.00058008 | 0.055398 |
| PKN2 | 4.96E-06 | 0.000421 | LATS2 | 0.00078832 | 0.060985 |
| CDK10 | 4.96E-06 | 0.000421 | TAOK1 | 0.00079823 | 0.060985 |
| CSK | 0.00032227 | 0.024621 | EIF2AK4 | 0.0015122 | 0.105028 |
| DLG1 | 0.0011453 | 0.079545 | MAPKAPK2 | 0.0020675 | 0.122669 |
| CIB4 | 0.001889 | 0.120265 | PRKRA | 0.0020873 | 0.122669 |
| MARK2 | 0.0020873 | 0.122669 | CDK4 | 0.0033962 | 0.185335 |
| TRIM28 | 0.0030392 | 0.162879 | MAPK6 | 0.0038821 | 0.197727 |
| PGM2L1 | 0.0031979 | 0.162879 | IRAK4 | 0.0048538 | 0.231771 |
| PHKA1 | 0.003892 | 0.185843 | CHEK2 | 0.0053199 | 0.239082 |
| ITPKA | 0.0048935 | 0.212753 | VRK3 | 0.0056967 | 0.241793 |
| CAMK2B | 0.0050125 | 0.212753 | AKT1 | 0.0076898 | 0.300645 |
| PRPF4B | 0.0055876 | 0.218147 | PFKFB1 | 0.0086616 | 0.300645 |
| CAD | 0.0060338 | 0.218147 | KSR1 | 0.0088400 | 0.300645 |

**B**

| DETROIT | | | | | |
|---|---|---|---|---|---|
| **id** | **neg\|p-value** | **neg\|fdr** | **id** | **pos\|p-value** | **pos\|fdr** |
| ATM | 4.96E-06 | 0.001263 | JAK1 | 4.96E-06 | 0.001263 |
| PRKDC | 4.96E-06 | 0.001263 | TYK2 | 4.96E-06 | 0.001263 |
| MAP3K7 | 4.96E-06 | 0.001263 | EXOSC10 | 4.96E-06 | 0.001263 |
| CAMK2B | 0.00097672 | 0.158333 | MKNK2 | 1.49E-05 | 0.002273 |
| NEK9 | 0.0010362 | 0.158333 | TAOK1 | 1.49E-05 | 0.002273 |
| MAP2K7 | 0.0022955 | 0.292298 | KSR1 | 2.48E-05 | 0.003157 |
| FGFR4 | 0.0031186 | 0.298972 | CRKL | 0.00040159 | 0.043831 |
| PEAK1 | 0.0040308 | 0.298972 | MAP4K4 | 0.0007784 | 0.074337 |
| FGFR3 | 0.0042192 | 0.298972 | ALDH18A1 | 0.0011056 | 0.093855 |
| ALPK2 | 0.0046952 | 0.298972 | HK2 | 0.0016609 | 0.121503 |
| LTK | 0.0051811 | 0.298972 | CDK15 | 0.0019881 | 0.121503 |
| CSF1R | 0.0053695 | 0.298972 | CSNK1G2 | 0.0020675 | 0.121503 |
| SCYL1 | 0.0053794 | 0.298972 | TTN | 0.0020675 | 0.121503 |
| PAK6 | 0.0054785 | 0.298972 | BCR | 0.0030591 | 0.166937 |
| PI4KB | 0.0065792 | 0.335101 | PKM | 0.0041796 | 0.212879 |
| LMTK3 | 0.0071246 | 0.338904 | ACVR2A | 0.0053298 | 0.254498 |
| MAP3K13 | 0.0075411 | 0.338904 | CSK | 0.0062718 | 0.281863 |
| SH3BP5L | 0.0082649 | 0.3508 | HUNK | 0.0070155 | 0.297769 |
| MAP2K6 | 0.013461 | 0.541268 | PIM1 | 0.0079079 | 0.30458 |
| MPP6 | 0.014849 | 0.567235 | NLK | 0.0085029 | 0.30458 |

**Supplemental Table 2: Sequences for control and JAK1-targeting gRNAs.** The top enriched gRNAs targeting JAK1 from the kinome-wide CRISPR screen were chosen from the optimized Brunello library. Indicated gRNAs were cloned into the lenti-Guide-Puro vector, and virally transduced into Cal27 and Detroit562 cells.

| **sgRNA Target** | **sgRNA Sequence** |
|---|---|
| Control | 5’ AAGATGAAAGGAAAGGCGTT 3’ |
| JAK1 (gRNA1) | 5’ GCCTAGACAGCACCGTAATG 3’ |
| JAK1 (gRNA2) | 5’ TTGATGACAAGATGTCCCTC 3’ |

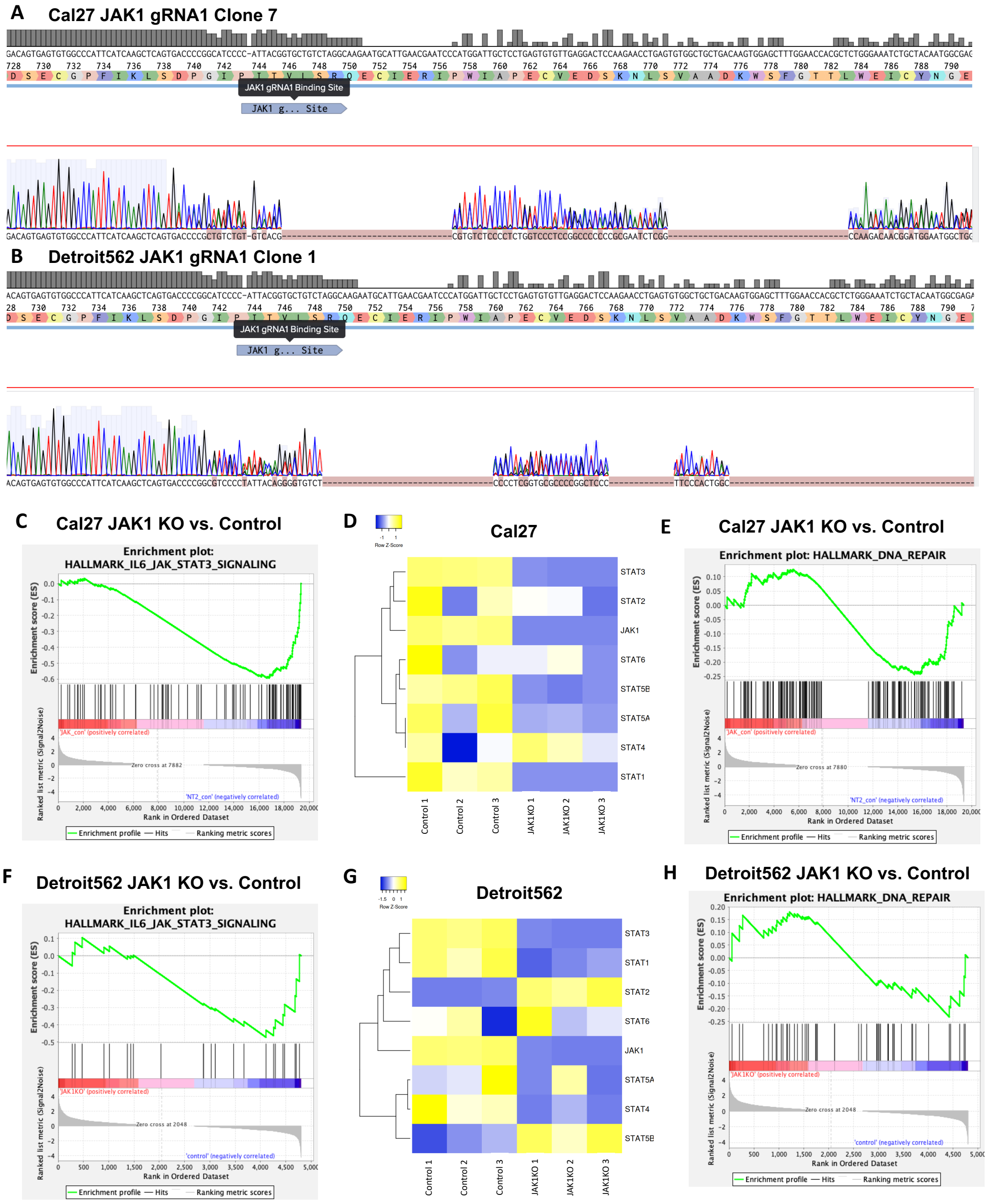


**Supplemental Figure 2: Confirmation of genomic mutation and loss of canonical JAK1 gene expression in chosen clones.** Genomic DNA from (A) Cal27 and (B) Detroit562 JAK1 clones was isolated. The genomic region in which the JAK1 gRNA1 binds was then amplified via PCR, and the product was sent for Sanger sequencing. Mutations for each clone are shown. (C-E) Cal27 and (F-H) Detroit562 control and JAK1 KO clones were sent for RNA sequencing to determine which genes were enriched and depleted in JAK1 KO cells compared to control cell lines. GSEA enrichment plots for the IL6-JAK-STAT3 hallmark gene set are shown for (C) Cal27 and (F) Detroit562 cell lines. This gene set was depleted in both JAK1 KO cell lines compared to control cells. Heatmaps for STAT gene transcripts are shown for (D) Cal27 and (G) Detroit562 cell lines. STAT transcripts were largely downregulated in JAK1 KO cell lines. There are no changes in DNA Repair transcriptomic signature between (E) Cal27 and (H) Detroit562 JAK1 KO and control cells.

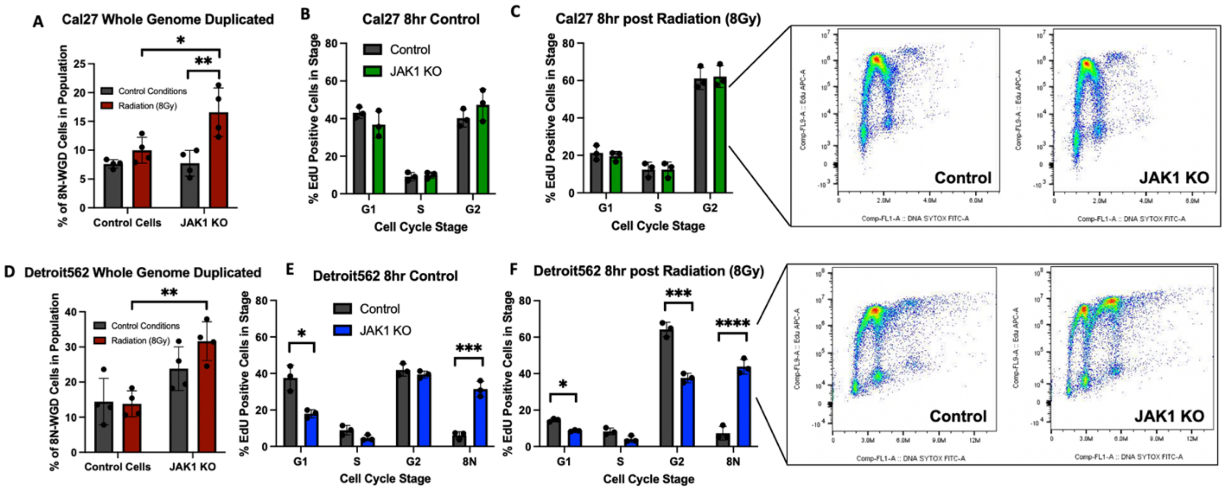


**Supplemental Figure 3: Presence of 8N-WGD cells and distribution of EdU-labeled population 8 hours following radiation or sham treatment.** (A) Cal27 and (D) Detroit562 cell lines were irradiated with 8 Gy or treated with a sham regimen. Twenty-four hours after treatment, cells were harvested and treated with propidium iodide to examine cell cycle distribution by DNA content via flow cytometry. Percentage of cells within the 8N-WGD population are shown. (B-C) Cal27 and (E-F) Detroit562 control and JAK1 KO cells were pulsed with 30uM of EdU for two hours to label the S phase population, irradiated with 8Gy or a sham regimen, and harvested 8-hours later. Cells were then treated with SYTOX green to determine DNA content distribution and analyzed via flow cytometry. The cell cycle distribution for the EdU-labaled population is shown.

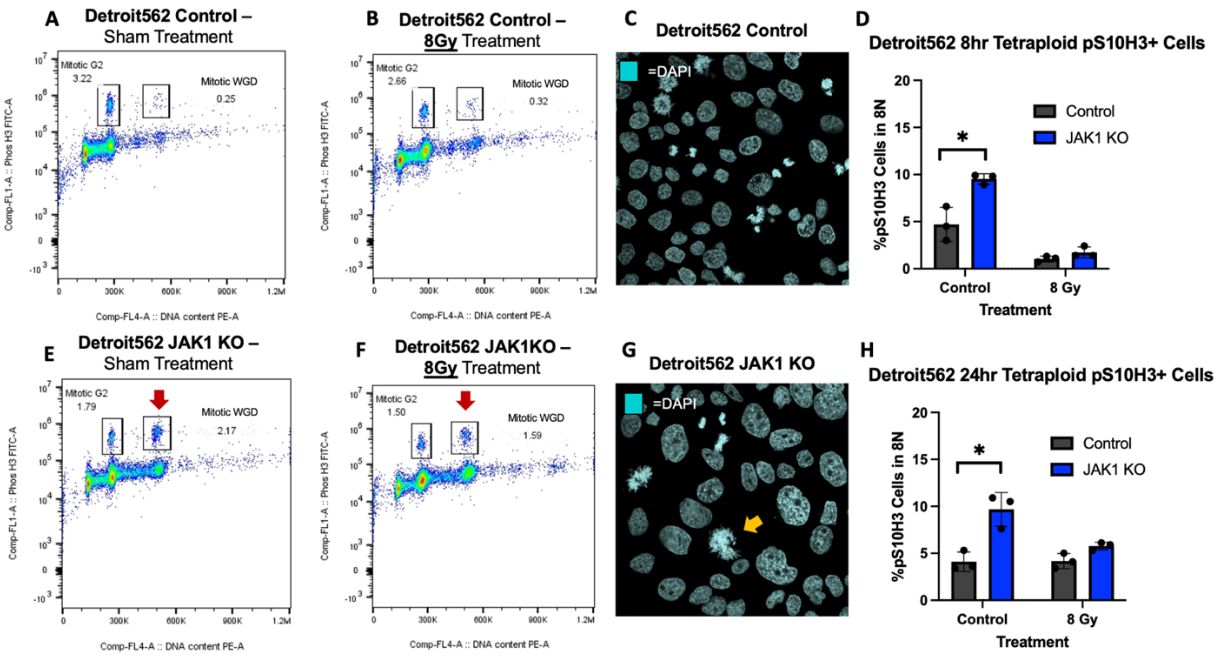


**Supplemental Figure 4: Detroit562 JAK1 KO cells have an 8N-WGD mitotic phosoho-Ser10 H3 population.** Detroit562 (A-B) control and (E-F) JAK1 KO cells were treated with 8Gy of radiation or a sham regimen and were harvested 24 hours later. Cells were stained for phosphor-Ser10 H3 (signal on Y axis) and DNA content using propidium-iodide (signal on X axis) and analyzed via flow cytometry. Red arrows indicate mitotic 8N-WGD population. Detroit562 (C) control and (G) JAK1 KO cells were seeded on glass coverslips, synchronized using a double thymidine block, released from the block, and fixed 12 hours after release. Cells were then incubated in a DAPI-Hoechst staining solution to visualize DNA. An 8N-WGD cell with condensed mitotic chromatin is indicated by the orange arrow. Detroit562 cells were irradiated or treated with a sham regimen and harvested (D) 8- or (H) 24-hours following treatment. Cells were stained for phosphor-Ser10 H3 and DNA content using propidium-iodide and analyzed via flow cytometry. The percentage of phosphor-Ser10 H3-positive events in the 8N-WGD population are shown. * indicates p< 0.0

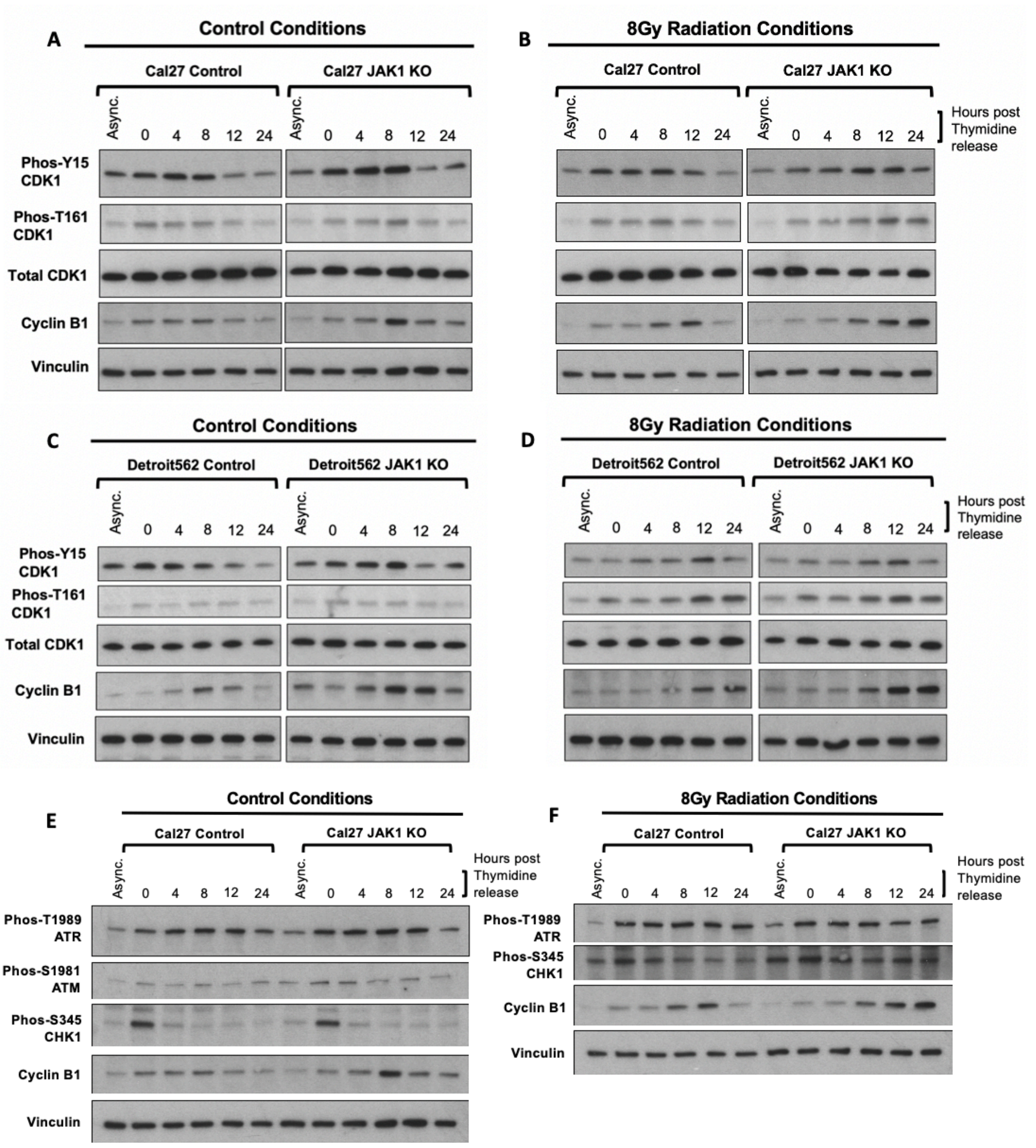


**Supplemental Figure 5: The canonical CDK1-dependent checkpoint nor hyperactivation of DNA damage signaling are responsible for JAK1 KO cell cycle phenotypes.** (A-B, E-F) Cal27 and (C-D) Detroit562 control and JAK1 KO cells were synchronized in G1 using a double thymidine block. Cells were then released from the block and immediate irradiated with 8Gy or treated with a sham regimen. Cells were then lysed at the indicated timepoints after thymidine release. Western blots were completed to examine changes in activation and accumulation of (A-D) CDK1 and Cyclin B1 as well as (E-F) ATM, ATR, and CHK1.

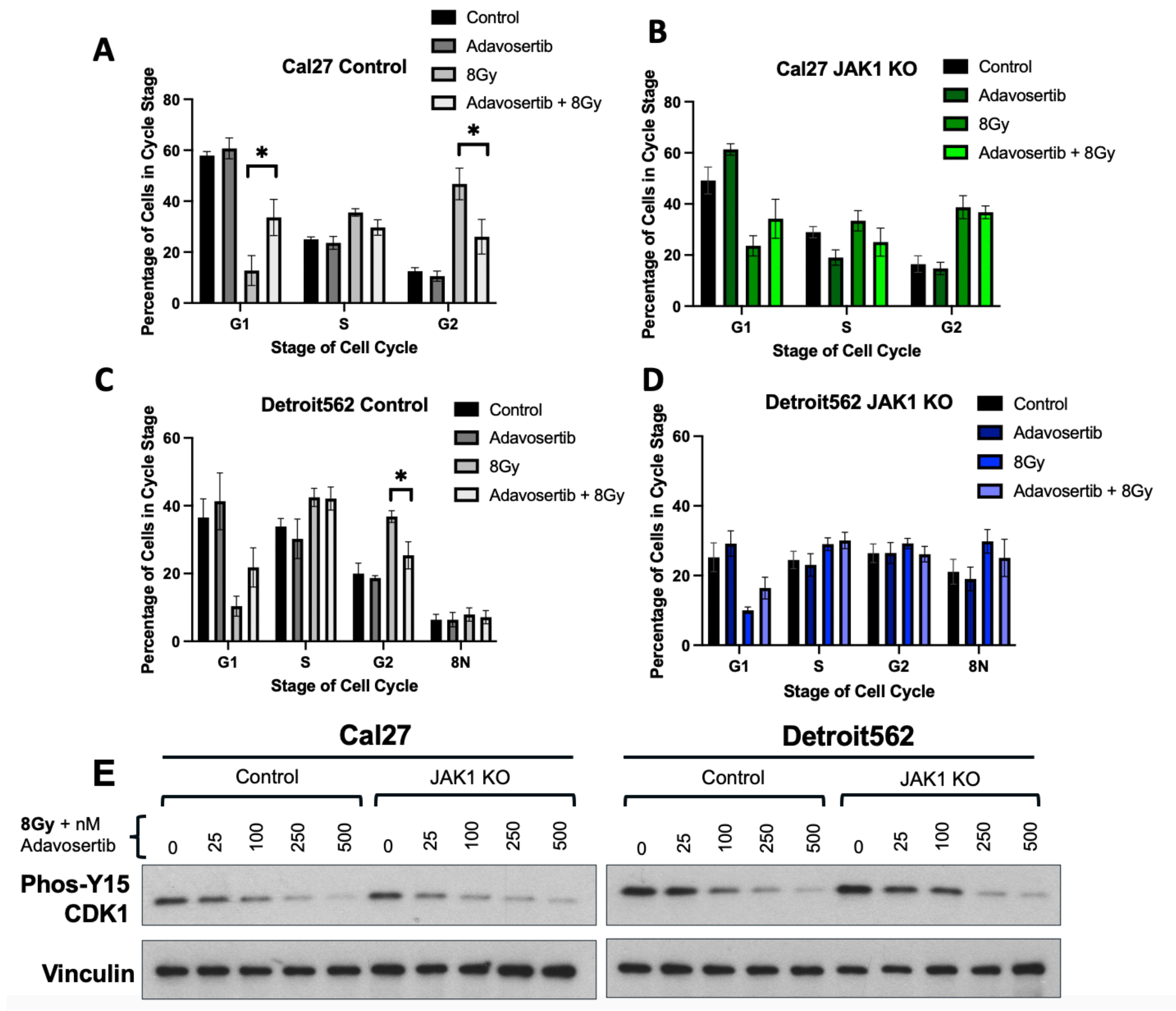


**Supplemental Figure 6: Inhibition of Wee1 is insufficient to abrogate JAK1 KO cell cycle changes despite sufficient dephosphorylation of CDK1 at Y15.** (A-B) Cal27 and (C-D) Detroit562 control and JAK1 KO cells were treated with 500nM of Adavosertib, 8Gy, or a combination of the two. Cell cycle was determined 8 hours later using propidium iodide staining and flow cytometry. (E) Cal27 and Detroit562 control and JAK1 KO cells were treated with 0-500nM of Adavosertib, and the phosphorylation status of Y-17 on CDK1 was determined. Loss of phosphorylation of Y-15 in CDK1 was observed upon Adavosertib treatment, consistent with successful inhibition of Wee1.

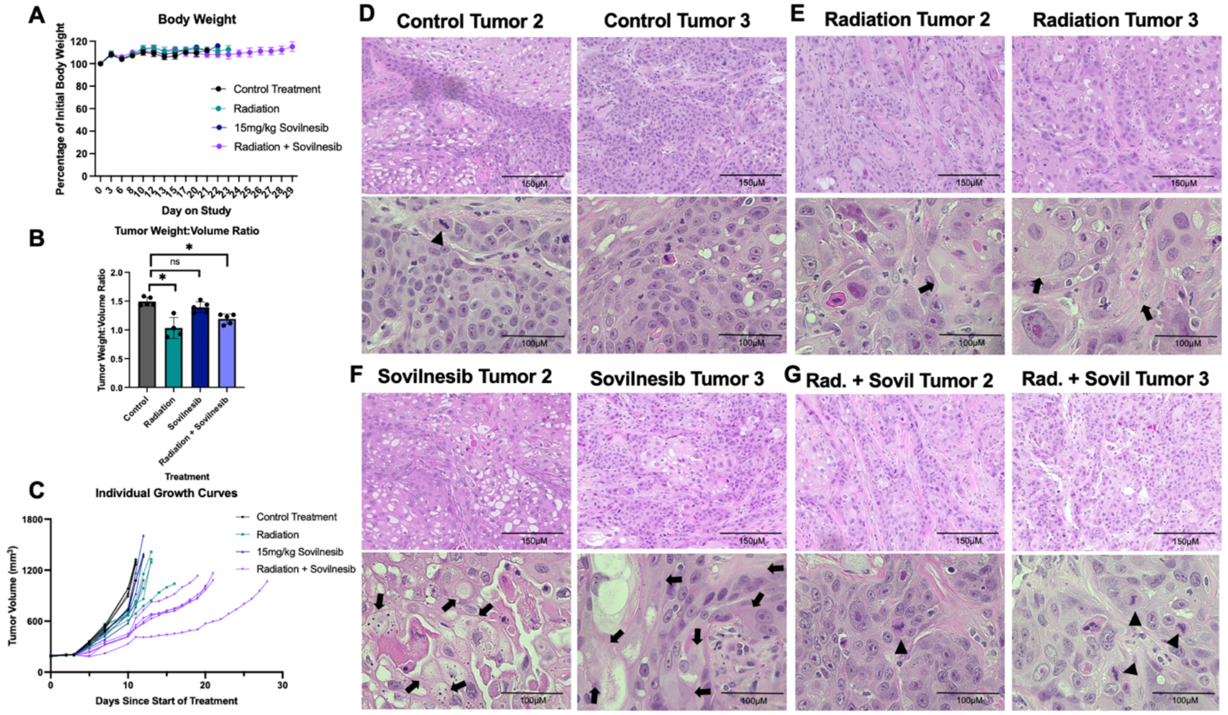


**Supplemental Figure 7: Kif18A inhibition sensitizes JAK1 KO cells to radiation treatment and causes changes in tumor architecture.** Detroit562 control JAK1 KO tumors were initiated in athymic nude mice. When tumors reached 200mm$^3$, mice were treated with a multifractionated radiation regimen of 2Gy per day, 15mg/kg of sovilnesib, a combined treatment of radiation and sovilnesib, or a sham regimen. (A) Mouse weight was monitored throughout the experiment and shown as a percentage initial body weight on day 1 of the study. sovilnesib, radiation, and combined treatment was well tolerated in the mice as little fluctuation in body weight was observed. (B) Tumor volume was determined using caliper measurements of flank tumors. Tumor weight was determined upon mouse sacrifice and tumor resection. Fluid filled pockets (of necrotic material) in the tumor were naturally emptied during this process. The ratio of tumor volume to tumor weight post dissection were determined for each tumor. (C) Individual tumor growth curves for each mouse in all treatment groups are shown. Three tumors per experimental group were chosen for haematoxylin and eosin (H&E) staining. Representative H&E images for the additional two tumors per group (tumor one shown in Figure 8) are shown for the (D) control, (E) radiation, (F) sovilnesib, and (G) radiation plus sovilnesib treatment groups. Black squares indicate areas consistent with tissue necrosis. Black triangles indicate areas containing mitotic cells.

### SUPPLEMENTAL MATERIALS AND METHODS

**Supplemental Methods Table 1: Drugs used in study and working concentrations.**

| Drug | Supplier | Catalog Number | Concentration |
|---|---|---|---|
| Abrocitinib | MedChem Express | HY-107429 | 100nM-1uM |
| Adavosertib | Gift from Dr. Barbara Burtness's Laboratory | N/A | 500nM |
| Sovilnesib | MedChem Express | HY-132840 | *In vitro*: 350nM<br>*In vivo*: 15 mg/kg |
| Thymidine | MedChem Express | HY-N1150 | 2mM |

**Supplemental Methods Table 2: List of primers used in the CRISPR Screen.**

| **P5 primers** | |
|---|---|
| **Name** | **Sequence** |
| P5 0 nt stagger | AATGATACGGCGACCACCGAGATCTACACTCTTTCCCTACACGACG CTCTTCCGATCTTTGTGGAAAGGACGAAACACCG |
| P5 1nt stagger | AATGATACGGCGACCACCGAGATCTACACTCTTTCCCTACACGACG CTCTTCCGATCTCTTGTGGAAAGGACGAAACACCG |
| P5 2 nt stagger | AATGATACGGCGACCACCGAGATCTACACTCTTTCCCTACACGACG CTCTTCCGATCTGCTTGTGGAAAGGACGAAACACCG |
| P5 3 nt stagger | AATGATACGGCGACCACCGAGATCTACACTCTTTCCCTACACGACG CTCTTCCGATCTAGCTTGTGGAAAGGACGAAACACCG |
| P5 4 nt stagger | AATGATACGGCGACCACCGAGATCTACACTCTTTCCCTACACGACG CTCTTCCGATCTCAACTTGTGGAAAGGACGAAACACCG |
| P5 6 nt stagger | AATGATACGGCGACCACCGAGATCTACACTCTTTCCCTACACGACG CTCTTCCGATCTTGCACCTTGTGGAAAGGACGAAACACCG |
| P5 7nt stagger | AATGATACGGCGACCACCGAGATCTACACTCTTTCCCTACACGACG CTCTTCCGATCTACGCAACTTGTGGAAAGGACGAAACACCG |
| P5 8nt stagger | AATGATACGGCGACCACCGAGATCTACACTCTTTCCCTACACGACG CTCTTCCGATCTGAAGACCCTTGTGGAAAGGACGAAACACCG |

**Supplemental Methods Table 3: List of Barcodes used in the CRISPR Screen.**

| Detroit562 P7 Barcodes | |
|---|---|
| **Name** | **Sequence** |
| P7 barcode B01 | CAAGCAGAAGACGGCATACGAGATATTGGATTGTGACTGGAGTTCAGACGTGTGCTCTTCCGATCTTCTACTATTCTTTCCCCTGCACTGT |
| P7 barcode B02 | CAAGCAGAAGACGGCATACGAGATATACTCGGGTGACTGGAGTTCAGACGTGTGCTCTTCCGATCTTCTACTATTCTTTCCCCTGCACTGT |
| P7 barcode B03 | CAAGCAGAAGACGGCATACGAGATTATGAGAAGTGACTGGAGTTCAGACGTGTGCTCTTCCGATCTTCTACTATTCTTTCCCCTGCACTGT |
| P7 barcode B04 | CAAGCAGAAGACGGCATACGAGATGCACAGTTGTGACTGGAGTTCAGACGTGTGCTCTTCCGATCTTCTACTATTCTTTCCCCTGCACTGT |
| P7 barcode B05 | CAAGCAGAAGACGGCATACGAGATCGTGGATTGTGACTGGAGTTCAGACGTGTGCTCTTCCGATCTTCTACTATTCTTTCCCCTGCACTGT |
| P7 barcode B06 | CAAGCAGAAGACGGCATACGAGATTAGTAGAAGTGACTGGAGTTCAGACGTGTGCTCTTCCGATCTTCTACTATTCTTTCCCCTGCACTGT |

**Cal27 P7 Barcodes**

| Name | Sequence |
|---|---|
| P7 barcode B07 | CAAGCAGAAGACGGCATACGAGATGCACGATTGTGACTGGAGTTCAGACGTGTGCTCTTCCGATCTTCTACTATTCTTTCCCCTGCACTGT |
| P7 barcode B08 | CAAGCAGAAGACGGCATACGAGATCGGTAGCCGTGACTGGAGTTCAGACGTGTGCTCTTCCGATCTTCTACTATTCTTTCCCCTGCACTGT |
| P7 barcode B09 | CAAGCAGAAGACGGCATACGAGATTAGTTCTTGTGACTGGAGTTCAGACGTGTGCTCTTCCGATCTTCTACTATTCTTTCCCCTGCACTGT |
| P7 barcode B10 | CAAGCAGAAGACGGCATACGAGATTACAAGTTGTGACTGGAGTTCAGACGTGTGCTCTTCCGATCTTCTACTATTCTTTCCCCTGCACTGT |
| P7 barcode B11 | CAAGCAGAAGACGGCATACGAGATATCACTGGGTGACTGGAGTTCAGACGTGTGCTCTTCCGATCTTCTACTATTCTTTCCCCTGCACTGT |

| | |
|---|---|
| P7 barcode B12 | CAAGCAGAAGACGGCATACGAGATCGCATCAAGTGACTGGAGTTCA GACGTGTGCTCTTCCGATCTTCTACTATTCTTTCCCCTGCACTGT |

**Supplemental Methods Table 4. List of antibodies used in this study.**

| **Antibody** | **Company** | **Catalog Number** | **Dilution** | **Solution** | **Application** |
|---|---|---|---|---|---|
| JAK1 | Cell Signaling | 3344S | 1:1,000 | 1% BSA | WB |
| GAPDH | Proteintech | 60004-1-Ig | 1:10,000 | 1% Milk | WB |
| Total STAT1 | Cell Signaling | 9172S | 1:1,000 | 1% BSA | WB |
| Phos-Y701-STAT1 | Cell Signaling | 9167S | 1:1,000 | 1% BSA | WB |
| Total STAT3 | Cell Signaling | 4904S | 1:1,000 | 1% BSA | WB |
| Phos-Y705-STAT3 | Cell Signaling | 9134S | 1:1,000 | 1% BSA | WB |
| IRF9 | Cell Signaling | 76684S | 1:1,000 | 1% BSA | WB |
| Vinculin | Cell Signaling | 13901S | 1:1,000 | 1% BSA | WB |
| Phos-T210-PLK1 | Cell Signaling | 9062S | 1:1,000 | 1% BSA | WB |
| Total PLK1 | Millipore | 05-844 | 1:5,000 | 1% Milk | WB |
| Phos-S46-TCTP | Cell Signaling | 5251S | 1:1,000 | 1% BSA | WB |
| Total TCTP | Cell Signaling | 5128S | 1:1,000 | 1% BSA | WB |
| PARP | Cell Signaling | 9532S | 1:1,000 | 1% Milk | WB |
| Total Aurora Kinase A | Cell Signaling | 14475S | 1:1,000 | 1% Milk | WB |

| | | | | | |
|---|---|---|---|---|---|
| Histone 3 | Cell Signaling | 4499S | 1:2,000 | 1% Milk | WB |
| BIM | Cell Signaling | 2933S | 1:1,000 | 1% BSA | WB |
| Phos-Y15-CDK1 | Cell Signaling | 4539S | 1:1,000 | 1% BSA | WB |
| Phos-T161-CDK1 | Cell Signaling | 9114S | 1:1,000 | 1% BSA | WB |
| Total CDK1 | Cell Signaling | 9116S | 1:1,000 | 1% BSA | WB |
| Cyclin B1 | Cell Signaling | 4138S | 1:1,000 | 1% BSA | WB |
| Phos-T1989-ATR | Cell Signaling | 30632S | 1:1,000 | 1% BSA | WB |
| Phos-T1981-ATM | Cell Signaling | 5883S | 1:1,000 | 1% BSA | WB |
| Phos-S345-CHK1 | Cell Signaling | 2360S | 1:1,000 | 1% Milk | WB |
| cGAS | Cell Signalling | 79978S | 1:200 | 10% horse serum, 10% goat serum, 80% 1XPBST | IF |
| Alexa Flour 488 goat anti-rabbit antibody | Invitrogen | A11034 | 1:500 | 10% horse serum, 10% goat serum, 80% 1XPBST | IF |
| Rad51 | EMD Millipore | PC130 | 1:300 | 10% horse serum, 10% goat serum, 80% 1XPBST | IF |

***Supplemental Flow Cytometry Methods:***

*Cell cycle analysis:* One million cells were seeded in a 10 cm dish and allowed to incubate for 48 hours. Cells were then treated and/or irradiated with 8 Gy. Twenty-four hours after radiation, cells were harvested via trypsinization, washed once in PBS, and fixed in 70% ice-cold ethanol. To prepare for flow cytometry, cells were washed in PBS and then incubated in an RNAse solution containing 200ug/ml Pure Link RNAse A (Invitrogen Cat#12091021) for 5 minutes at room temperature. Propidium Iodide (Sigma Cat#P4170) was then added to the cells in RNAse solution to achieve a final concentration of 40ug/ml Propidium Iodide and 40ug/ml Pure Link RNAse A. Samples were incubated for an additional 20 minutes in the dark at room temperature before proceeding to flow cytometry.

Samples were analyzed on a CytoFLEX LX instrument (Beckman Coulter) in the PE channel. On the instrument, cell nuclei were identified in a PE-A verses events histogram, and 20,000 nuclear events were collected per sample. All data was subsequently analyzed using FlowJo software version 10.8.1 (Becton Dickinson & Company). Cellular events were gated out from debris using SSC-H vs. FSC-H scatter plot. Doublet discrimination was then performed by comparing FSC-H vs. FSC-A. Single cells were then plotted in a PE-A vs. event number histogram. Cell cycle distribution was determined using the cell cycle tool in Flow Jo software.

*Phos-Ser10 H3 mitosis assays:* Cells were seeded and irradiated as described in "cell cycle analysis." Eight or 24 hours after radiation treatment, cells were harvested via trypsinization, washed once in PBS, and fixed in 70% ice cold ethanol overnight. Cells were then washed with PBS and permeabilized in 0.25% PBS-triton on ice for 15 minutes. Cells were then washed twice in 1% BSA in PBS, and then incubated in 1:30 dilution of Anti-phospho H3 (Ser10) antibody (clone 3H10 Alexa Fluor® 488 Sigma Aldrich, Cat# FCMAB104A4) in 1% BSA in PBS for one hour at room temperature. Cells were washed in 1% BSA in PBS and then resuspended in nuclear staining solution containing 40ug/ml Propidium Iodide and 40ug/ml Pure Link RNAse A (as previously described). Compensation matrixes were generated using single-stained samples, and all samples were analyzed on a CytoFLEX LX instrument. Twenty-thousand nuclear events were collected per sample.

All data was analyzed in Flow Jo software. Cellular event isolation and doublet discrimination were performed as previously described. Bivariate scatter plots comparing PE-A vs. A488-A were generated, and actively mitotic populations were gated accordingly. Cell cycle distribution was determined using the cell cycle tool within Flow Jo as previously described[17]. Percentage of actively mitotic populations were determined by dividing the amount of actively mitotic events by the amount of events in the G2 or hyperdiploid cell cycle population.

*EdU assays:* One million cells were seeded in 10cm dishes and incubated for 48 hours. Cells were then plused with 30 uM of EdU for two hours. The cell media containing EdU was then removed, plates were washed with PBS, and new cell media was added. Cells were irradiated with 8Gy or treated with a sham regimen, and allowed to incubate for either 8 or 24 hours prior to harvest. After incubation, cells were tyrpsonized, counted, and 2 million cells were used as an input for the Invitrogen Click-iT EdU Flow Cytometry Assay Kit (Cat# C10424) according to manufacturers instructions. SYTOX Green nucleic acid stain (Invitrogen Cat#S7020) was used as a DNA stain. Cell distribution as then determined via flow cytometry using a CytoFLEX LX instrument.

*Double thymidine synchronization:* Two million cells were seeded in a 10cm dish and incubated for 24 hours. Cells were then treated with 2mM of thymidine (Med Chem Express Cat# HY-N1150) for 17 hours. Cells were then rinsed with PBS and incubated in regular media for 8 hours. Thymidine was then added to a final concentration of 2mM for and additional 16 hours. Cells were released from thymidine by rinsing with PBS and then adding regular media. For thymidine-synchronization and radiation experiments, cells were released from thymidine and immediately irradiated as previously described or treated with a sham regimen. For cell cycle analysis, cells were harvested via trypsinization and analyzed as previously described. For western blot analysis, cells were lysed directly in the plate and analyzed as previously described.

***Supplemental Sovilnesib clonogenic methods:***

For multifractionated radiation clonogenics with sovilnesib, 1 million Cal27 cells were seeded in 10cm dishes. Twenty-four hours after seeding, dishes were pre-treated with 350nM of sovilnesib or vehicle for one hour. Plates were then irradiated with 2Gy every 24 hours to achieve the indicated total radiation doses. After multifractionated radiation treatment, cells were re-seeded at a density of 1000-6000 cells per well in a 6 well plate. All clonogenics were incubated for 14 days after re-seeding and then stained 0.25% crystal violet and 80% methanol. Colonies with greater than 50 cells were hand counted. The surviving fraction of each sample was calculated as the ratio between the number of colonies counted divided by number of cells seeded and the plating efficiency, thus normalizing for plating efficiency differences with treatment. Clonogenic survival differences for each treatment were compared using survival curves generated from the linear quadratic equation as previously described.
